# PfPPM2 signalling regulates asexual division and sexual conversion of human malaria parasite *Plasmodium falciparum*

**DOI:** 10.1101/2024.10.01.615984

**Authors:** Akanksha Rawat, Neelam Antil, Meenakshi, Bhagyashree Deshmukh, Akhila Balakrishna Rai, Narendra Kumar, T.S. Keshava Prasad, Krishanpal Karmodiya, Pushkar Sharma

## Abstract

Malaria parasite transits through distinct developmental stages during its life cycle in the human and mosquito host, which includes unique asynchronous division in the erythrocytes. The switch from its asexual stage to sexual forms, which is critical for disease transmission, is intricately regulated but signalling pathways involved in this process have remained unknown. In the present study, we report a novel signalling pathway involving Protein Phosphatase PfPPM2, which regulates asexual division of the parasite as well as its conversion to sexual forms. Phosphoproteomics revealed that PfPPM2 may regulate the phosphorylation of key proteins involved in chromatin remodelling and protein translation. One of the key PfPPM2-targets that emerged from these studies was Heterochromatin Protein 1 (HP1), a regulator of heritable gene silencing which contributes to both mitotic proliferation as well as sexual commitment of the parasite. We demonstrate that PfPPM2 promotes sexual conversion by regulating the interaction between HP1, H3K9me3 and chromatin and it achieves this by dephosphorylating S33 of HP1. Regulation of HP1 and Histone H3 by PfPPM2 may also contribute to division. In addition, PfPPM2 also regulates protein synthesis in the parasite by repressing the phosphorylation of initiation factor eIF2α, which is likely to contribute to parasite division and possibly sexual differentiation.

## Introduction

Malaria continues to be a major public health challenge as it causes ∼200 million infections and more than 600,000 deaths each year (World Health Oganization, 2022). *Plasmodium,* the causative agent of malaria, has an unusual life cycle involving morphologically distinct developmental stages. It undergoes asexual division in hepatocytes in the liver followed by asynchronous intraerythrocytic multiplication in the vertebrate host. In each cycle, a small fraction of the asexual parasites convert to nonreplicative sexual gametocytes, which upon ingestion by the mosquito initiate the sexual development (Cowman et al, 2016). Reversible protein phosphorylation mediated by protein kinases and phosphatases are major constituent of signalling pathways and are involved in the development of most organisms. The fact that several of the ∼85 kinases and ∼27 phosphatases coded by *Plasmodium* genome are indispensable for parasite growth (Adderley & Doerig, 2022; Guttery et al, 2014), reiterates their importance for parasite biology and also suggests that at least some of these may serve as drug targets. The function of most protein phosphatases remains unknown and the organization of parasite signalling pathways is poorly understood.

Present study relate to protein phosphatase PfPPM2 from human malaria parasite *Plasmodium falciparum*, which belongs to PPM2 family of phosphatases (Guttery et al, 2014). It is refractory to gene disruption in *P. falciparum*, which indicates that it is indispensable for asexual blood stage development of the parasite (Mamoun & Goldberg, 2001; Zhang et al, 2018). Interestingly, its *P.berghei* homologue PbPPM2 was found to be dispensable for the asexual development in the rodent parasite but plays a role in gametocyte sex allocation and ookinete differentiation (Guttery et al, 2014), however, the underlying mechanisms have remained unclear. We demonstrate that PfPPM2 is involved in asexual division of *Plasmodium falciparum* and it also regulates sexual differentiation of this parasite. Detailed investigations revealed that it may control these processes by regulating the phosphorylation of key proteins involved in chromatin remodelling as well as protein translation. Heterochromatin Protein 1 (HP1) was identified as one of its key targets, which was relevant for both asexual division and sexual differentiation of the parasite. In addition, PfPPM2 regulates protein synthesis by regulating eIF2α phosphorylation by eIF2α-kinase PfPK4.

## Results

### PfPPM2 regulates asexual division of P. falciparum

PfPPM2 belongs to Metal Dependent Protein Phosphatase (PPM) family,which is regulated by Mg^2+^ and Mn^2+^ (Wilkes & Doerig, 2008). Interestingly, the catalytic domain of PfPPM2 is interrupted by a linker and it also has a putative N-terminal myristoylation signal (Guttery et al, 2014) Given that PfPPM2 is refractory to gene disruption (Zhang et al, 2018), an inducible/conditional knockdown approach involving autocatalytic glmS ribozyme, which can be activated by using glucosamine (GlcN), was used (Prommana et al). To this end, glmS ribozyme was introduced into the PfPPM2 3’-UTR along with a 3xHA tag (Fig. 1A, Supp. Fig. 1) in 3D7 (PfPPM2-HA-glmS^3D7^) as well as NF54 (PfPPM2-HA-glmS^NF54^) background. The treatment of PfPPM2-HA-glmS parasites with GlcN effectively depleted PfPPM2 protein (Fig. 1B). Parasite growth rate assays were performed in which glucosamine (GlcN) was added at early ring stages (cycle 0) and parasite development was monitored periodically for at least two subsequent cycles. A marked decrease in parasitemia was observed at the end of the first cycle, which was even more accentuated in the second cycle (Fig. 1C). Presence of GlcN itself did not affect the asexual proliferation in either of 3D7 or NF54 parasite lines (Supp. Fig. S2). The analysis of Giemsa-stained thin blood smears did not reveal any obvious morphological changes in the parasite, which was also indicated by similar distribution of various developmental stages in PfPPM2-depleted parasites (Supp. Fig. S3A) Further, no significant defects in invasion were observed (Supp. Fig. S3C). Strikingly, PPM2 depletion led to a significant reduction in the number of merozoites formed per schizont (Fig. 1D). These data were suggestive of a role of PfPPM2 in parasite division. Immunofluorescence assays revealed that PfPPM2 is present in punctate structures near the parasite nucleus during schizogony, particularly in mid to late schizonts, with some presence in the cytoplasm. However, in trophozoites, PfPPM2 it was more cytoplasmic and did not co-localize with MTOC components such as CPs (Fig. 1E). Further, IFAs were performed to look at the division of centriolar plaque (CP)-which is analogous to the centrosomes in mammalian cells and also represents the microtubule organization centre (MTOC) of the parasite and spindle formation (Arnot et al, 2011; Gerald et al, 2011). The mitotic spindle was also visualized by staining with anti-α-tubulin antibody (Liffner et al, 2023). PfPPM2 depletion revealed parasites with diffused tubulin staining indicative of loss of spindles (Fig. 1F). In addition, there was a modest but significant defect in CP duplication (Fig. 1G). In addition, the formation of inner membrane complex (IMC) assessed by IMC1g staining (Kono et al, 2012), which is formed during division and speculated to support this process, was also disrupted and may be a consequence of arrested nuclear division (Fig. S3D). Collectively, these results indicated a role of PfPPM2 in asexual division of the parasite, which may be due to its direct or indirect involvement in the above-mentioned processes (discussed later in detail).

**Figure 1:**
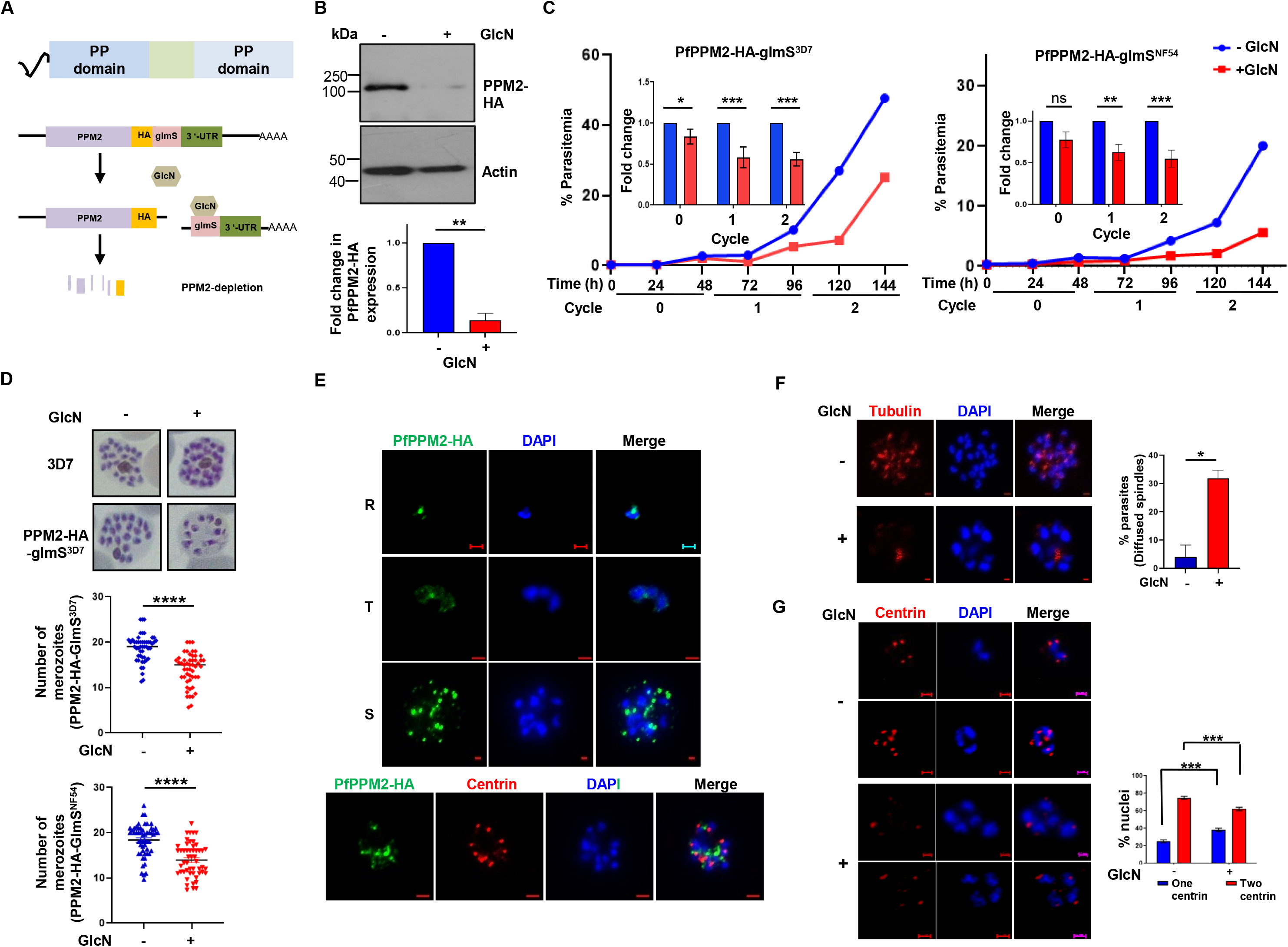
PfPPM2 regulates asexual division of *Plasmodium falciparum*. A. PfPPM2 has an insert between its PP domain and also has a putative N-terminal myristylation signal (Guttery et al, 2014; Wright et al, 2014). In order to achieve conditional knockdown (cKD) of *PfPPM2*, a ribozyme glmS was introduced at its 3’end along with a HA-tag in 3D7 as well as NF54 background (Supp. Fig. S1). The addition of glucosamine activates the ribozyme and will results in its cleavage and degradation of the mRNA causing depletion of PfPPM2 protein. B. Western blotting of PfPPM2-HA-glmS^3D7^ parasites, which were either left untreated or treated with GlcN (cycle 0, rings) and parasite lysates were prepared (schizonts, cycle 1) using anti-HA antibody, which caused effective depletion of PfPPM2-HA. β-actin antibody was also used to probe the blots. *Bottom Panel*, Fold change in PfPPM2 depletion was determined by densitometry of PfPPM2 bands in the Western blots, which was normalized with respect to actin (SEM ± SE, n=3, t-test, **P<0.01). C. PfPPM2-HA-glmS^3D7/NF54^ parasites were synchronized and ring stage parasites were used for setting up growth rate assay in the presence or absence of GlcN. Parasite growth was assessed after each cycle by performing flow cytometry (right panel) as well as analysis by counting parasites from Giemsa-stained smears (left panel). *Inset*, fold change in parasitemia upon GlcN treatment was determined (SEM ± SE, n=4, ANOVA, *P<0.05,**P<0.01, ***P<0.001). D. PfPPM2-HA-glmS^3D7/NF54^ parasites were treated with GlcN as described in panel B. Thin blood smears were made at ∼40-44 hpi in cycle 1. The number of nuclear centers/merozoites were counted from Giemsa-stained thin blood smears (SEM ± SE, n=3, paired t-test, ****P<0.0001). E. PfPPM2-HA-glmS^3D7^ parasites were synchronized and IFA was performed on Rings (R), trophozoites (T) and Schizonts (S) stages using anti-HA antibody. The localization of PfPPM2-HA is prominent during schizogony, and is localized close to the nucleus and/or in its close proximity (DAPI). *Bottom Panel,* IFA image showing mature schizonts co-stained with antibody against centrin. F and G. PfPPM2-HA-glmS^3D7^ parasites were treated with GlcN as described in panel B and C. Thin blood smears were made ∼40-44 hpi in cycle 1. IFA was performed using anti-α-tubulin (F) or Centrin-1 (G) antibodies. Scale bar =1 μm. *Right Panel*, parasites with defects in tubulin staining (F) or in the number of centrin puncta resembling CPs were counted from IFA images (F, Mean ± SEM, t-test, N=3, * p<0.05), (G, Mean ± SEM, ANOVA, N=3,, *** p<0.001, P > 0.05, ns-non-significant).

### Identification of PfPPM2 targets by phosphoproteomics

Having established the role of PfPPM2 in parasite division, efforts were made to assess the mechanisms via which it may regulate this process, for which it was important to identify its targets or substrates in the parasite. For this purpose, a comparative phosphoproteomics approach was used (Supp. Fig. S4) to compare the phosphoproteome of parasites after PfPPM2-depletion. Since a major defect was found in parasite division, schizonts (∼38-40 hpi, 1st cycle) that were left untreated or treated with GlcN were used for this purpose (Supp. Fig. S4). Overall, 95 phosphosites on 61 proteins exhibited enhanced phosphorylation upon phosphatase depletion (Fig. 2A, Supp Data Set S1.2). In addition, 35 phosphosites on 27 proteins were identified as hypophosphorylated in the PfPPM2-depleted parasites, which may be due to direct or indirect effects exerted by PfPPM2 on a protein kinase or by negatively regulating other protein phosphatases.

**Figure 2.**
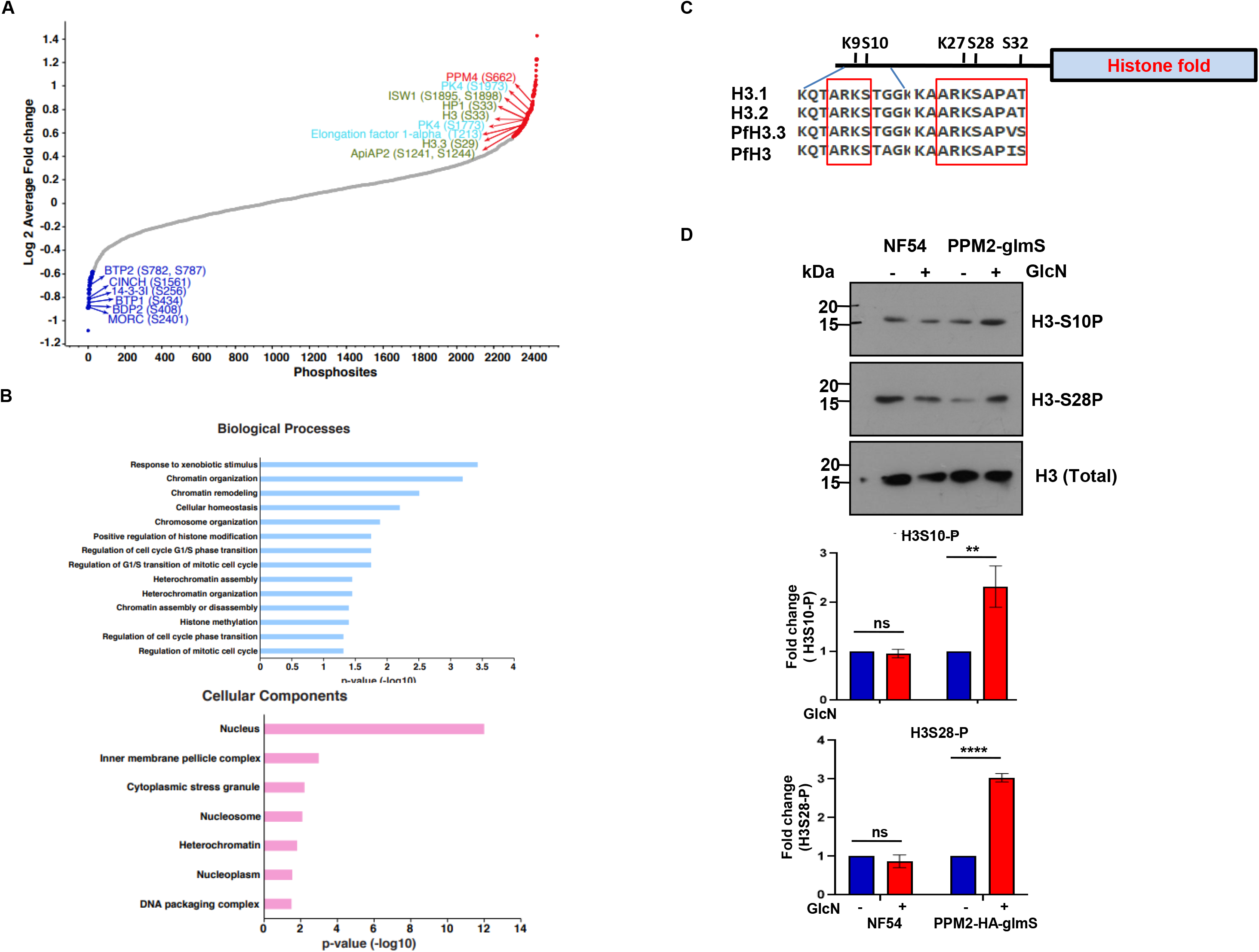
PfPPM2 regulates the phosphorylation of key parasite proteins involved in chromatin remodelling and translation. A. PfPPM2-HA-glmS^3D7^ parasites were treated with GlcN as described in Fig. 1B and C to perform comparative phosphoproteomic and proteomic analyses as indicated in the schematic (Supp. Fig. S4). The Scatter-curve represents fold-change ratios of identified phosphopeptides upon GlcN treatment (Supplementary Data Set S1.1). Some of the significantly altered phosphorylation sites belonging to key proteins relevant for protein synthesis (blue) or chromatin organisation (green) (Supplementary Data Set 1.2, 1.3) are indicated. B. Gene Ontology enrichment of proteins that exhibited significant changes in phosphorylation upon PfPPM2 depletion (Supplementary Datset 1.3,1.4). C. Sequence comparison of N-terminal tail of human histone H3 variants with *P. falciparum* Histone H3 and H3.3. Key motifs containing regulatory sites indicated by red square and critical residues that are modified by phosphorylation (S10, S28, S32), methylation (K9) and acetylation (K9, K27) are indicated. D. Western blot analysis using specific antibodies against pS10-H3, pS28-H3 performed on NF54 and PfPPM2-HA-glmS^NF54^ lines cultured in the absence and presence of GlcN. *Lower Panel*, densitometry analysis was performed and fold change in GlcN-treated parasites compared to untreated parasites was determined (Mean ± SEM, ANOVA, N=3, ** p<0.01, **** p<0.0001, P > 0.05, ns-non-significant).

The differentially phosphorylated proteins in PfPPM2 depleted parasites fell into three major family of proteins (Fig. 2A, Supp Dataset S1): chromatin remodelling, assembly and related functions (histone H3 and H3 variant 3.3, Heterochromatin Protein 1, ISWI chromatin remodelling complex ATPase); protein translation (eukaryotic translation initiation factor 2-alpha kinase, elongation factor 1-α,); IMC formation [(basal complex transmembrane protein 1/2 (BTP1/2)], CINCH (Table S1). Interestingly, each of these processes can affect parasite division, which as mentioned above is regulated by PfPPM2. For instance, both protein synthesis (Polymenis & Aramayo, 2015) and chromatin organization are critical for cell division. Epigenetic factors involved in chromatin organization and remodelling are known to be critical for the tight regulation and timely expression of genes in *P. falciparum*. IMC formation has been implicated in the division of Apicomplexan parasites *Toxoplasma* and *Plasmodium* (Beck et al, 2010; Kono et al, 2012). As mentioned above, PfPPM2 depletion impaired IMC biogenesis (Fig. S3D), which may be due to altered phosphorylation of the above-mentioned proteins. Since PfPPM2 was not localized to the IMC, it is possible that it may not directly regulate IMC biogenesis, instead, it may be an outcome of arrest in parasite division.

### PfPPM2 may regulate epigenetic regulators and components of translation machinery

Since both phosphoproteomics (Fig. 2A, Table S1, Dataset S1) and transcriptomics (Supp. Fig. S9, discussed below) indicated that PfPPM2 may regulate genes/proteins involved in translation, its role in protein synthesis was assessed. Interestingly, phosphoproteomics data suggested that PfPPM2 may regulate the phosphorylation of eIF2α-kinase PfPK4 as two residues (S1773, S1973) were found to be hyperphophorylated upon PfPPM2 depletion (Fig. 2A, Table S1, Supp. Fig. S5A). In the case of malaria parasite, protein kinase PK4 has been implicated in eIF2α phosphorylation at S59, which prevents protein synthesis (Supp. Fig. S5B) (Zhang et al, 2017). Western blotting was performed using anti-phospho-eIF2α antibody, which recognizes the complementary site (S59) on its *Plasmodium* homologue (Zhang et al, 2017; Zhang et al, 2012). PfPPM2 depletion resulted in a dramatic increase in eIF2α phosphorylation (Supp. Fig. S5C) suggesting that PfPPM2 normally suppresses this during asexual development. Given that phosphorylation of eIF2α at this site is critical for the regulation of protein translation in several organisms including *Plasmodium* (Zhang et al, 2017; Zhang et al, 2012), changes in protein synthesis were assessed. To this end, puromycin incorporation assays were performed to assess the status of protein synthesis (McLean & Jacobs-Lorena, 2017; Thommen et al, 2023). Strikingly, PfPPM2 depletion caused a marked reduction in puromycin incorporation, which was suggestive of impaired protein synthesis (Supp. Fig. S5D). Therefore, PfPPM2 may negatively regulate PK4 activity and/or dephosphorylate eIF2α at this site to promote protein synthesis.

One of the highlights of phosphoproteomic studies was the aberrant phosphorylation of several proteins involved in chromatin remodelling, which included HP1, Histone H3 and its variant H3.3 in PfPPM2-depleted parasites (Fig. 2A, Table S1, Dataset S1). Histone H3 and its variant H3.3 were found to be hyperphopshorylated at S33, S29 (corresponding to S28 on histone H3), respectively. Since S29 is equivalent to S28 of histone H3 and H3.3 homlogues of most species, it will be hereafter referred to as S28 (Dastidar et al, 2013). In the case of mammalian cells, phosphorylation of S28 is very well known for its role in mitosis and can act as a distinctive mark for dividing cells (Gil & Vagnarelli, 2019; Komar & Juszczynski, 2020; Sawicka & Seiser, 2012). Although a phosphopeptide with phospho-S10 H3 was not detected, it is also known to be critical for cell division. Since these sites are very well conserved in PfHistone H3 and its variant H3.3 (Fig. 2C), it is very likely that the phosphorylation of these sites is important for a similar function in the malaria parasite. Therefore, the role of PfPPM2 in S10 and S28 H3 phosphorylation was assessed using specfic commercial phospho-S10/S28 antibodies. Strikingly, the phosphorylation of both these sites was significantly enhanced upon PfPPM2 depletion (Fig. 2D), which confirmed findings from the phosphoproteomics analysis (Table S1, Data Set S1) and implicated a possible direct or indirect role of PfPPM2 in the dephosphorylation of these sites. Previous studies performed on mammalian cells have indicated that dephosphorylation of these and other sites on Histone H3 is critical before cells exit the M-phase failing which they undergo arrest (Sawicka & Seiser, 2012). Therefore, it is possible that PfPPM2 mediated dephosphorylation of Histone H3 contributes to parasite division in concert with other chromatin remodelling proteins indicated above.

Since the aberrant phosphorylation of histone H3 and other epigenetic regulators is likely to impact gene expression, it was pertinent to assess changes in parasite transcriptome upon PfPPM2 depletion. RNA-sequencing (RNA-Seq) analysis of PfPPM2-HA-glmS^Nf54^ parasites at schizont stage revealed several genes that exhibited altered expression upon PfPPM2 depletion (Supp. Fig. S9A and Supplementary Data Set S3). Gene Set Enrichment Analysis (GSEA) using ClusterProfiler (Yu et al, 2012) highlighted significant enrichment of Gene Ontology (GO) terms involved in translation as well as cell division (Supplementary Fig. S9B). For instance, two centrins(Centrin 4 and Centrin 1), PCNA and MCM3- were downregulated in PfPPM2-depleted parasites (Supp. Fig. S9, Data Set S3), which fits in well with defects in MTOC/CP duplication and division of these parasites (Fig. 1G). Additionally, several genes involved in translation showed altered expression (Supp. Fig. S9). This is particularly noteworthy as phosphoproteomics also revealed that numerous key proteins associated with translation displayed changes in phosphorylation upon PfPPM2 depletion (Fig. 2A,B Table S1). Importantly, as described above, PfPPM2 indeed regulates protein synthesis by suppressing the phosphorylation of eIF2α. Given that both chromatin remodelling and protein synthesis regulate cell division, these results provided first insights to the mechanisms by which PfPPM2 may regulate parasite division.

### PfPPM2 regulates HP1, which is essential for sexual differentiation and asexual development

PfHP1 is a heterochromatic protein that maintains heterochromatin structure and plays roles in var gene silencing, sexual differentiation, and asexual development of the parasite (Brancucci et al, 2014; Filarsky et al, 2018; Flueck et al, 2009; Perez-Toledo et al, 2009). Phosphoproteomics analysis revealed that S33 present in the chromodomain (CD) of PfHP1 was hyperphosphorylated upon PfPPM2 depletion. While this phosphorylation site is conserved in human HP1 homologues, it is replaced by an aspartic acid residue in *S. pombe* Swi6(Fig. 3A). Western blot using an custom antibody, generated against a phosphopeptide containing the sequence encompassing phospho-S33 (pS33-HP1), also revealed that phosphorylation of S33 was significantly enhanced upon PfPPM2 depletion, supporting phosphoproteomics data and suggesting that PfPPM2 may dephosphorylate this site on HP1 (Fig.3B).

**Figure 3:**
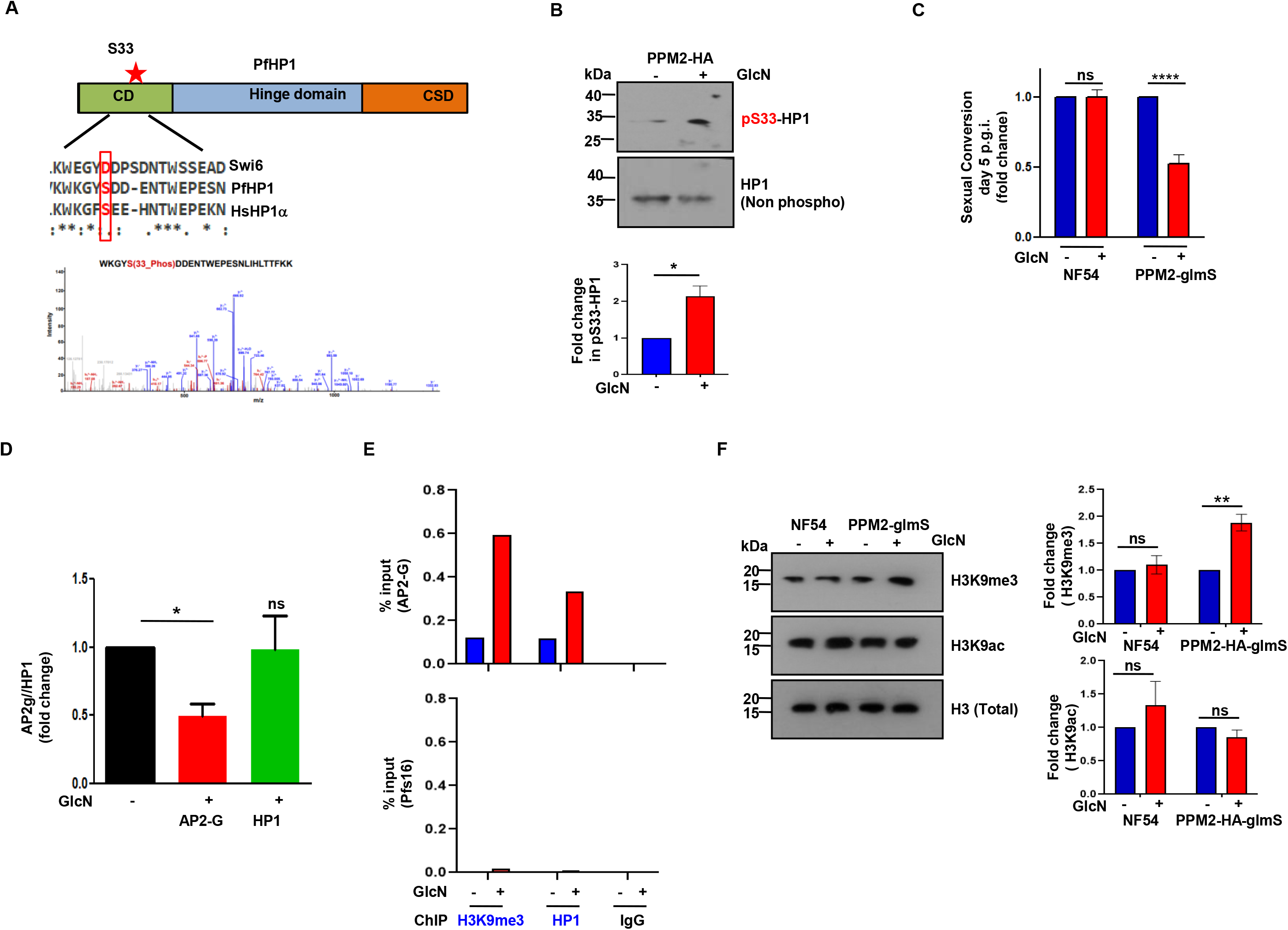
PfPPM2 regulates HP1 phosphorylation and sexual conversion of the parasite. A. PfPPM2 depletion causes hyperphosphorylation of S33 (*) present in its chromodomain (CD). S33 seems to be conserved in human HP1α and HP1β (not shown here) and is replaced by a D in *S. pombe* homologue swi6. *Bottom panel*, MS/MS spectra for peptide corresponding to pS33-HP1.B. Western blotting of lysates from PfPPM2-HA-glmS^NF54^ parasites, which were either left untreated or treated with GlcN using antibody against pS33-HP1 (see methods for details) or its phosphorylation-independent version. *Bottom Panel,* Fold change in HP1-S33 phosphorylation upon PfPPM2 depletion in experiments described in the upper panel was determined by densitometry (SEM ± SE, n=3, t-test, *P<0.05). C. PfPPM2-HA-glmS^NF54^or NF54 (control) parasites were cultured in the presence or absence of GlcN and gametocyte formation was induced as described in Experimental Procedures. Fold change in sexual conversion rates for both parasite lines upon GlcN treatment was determined at day 5 p.g.i. (Mean ± SEM, ANOVA, N=3, **** p<0.0001; ns- not significant). D. Gametocyte formation was induced in GlcN-treated or untreated PfPPM2-HA-glmS^NF54^ parasite. qRT-PCR was performed after 24h of gametocyte induction to compare the expression of AP2-G and HP1 (Mean ± SEM, ANOVA, N=3; *p<0.1, ns p>0.05). E. ChIP-qpCR was performed to determine PfHP1 and H3K9me3 occupancy at AP2-G or Pfs16 loci in PfPPM2-HA-glmS^NF54^ parasites which were either left untreated or treated with GlcN. IgG antibody was used for ChIP as a negative control. A representative of three independent replicates is shown. F. Western blotting of lysates from PfPPM2-HA-glmS^NF54^ or NF54 parasites, which were either left untreated or treated with GlcN using antibodies against H3K9me3, H3K9ac or total H3. *Right Panel,* Fold change in H3K9me3 upon PfPPM2 depletion in experiments described in the above panel was determined by densitometry (SEM ± SE, ANOVA n=3, NF54, **P<0.01, ns p>0.05).

PfHP1 is known to regulate gametocytogenesis or sexual differentiation (Brancucci et al, 2014). Given PfPPM2 regulates its dephosphoryation (Fig. 2A, Table S1, Dataset S1), we examined if PfPPM2 had a role to play in sexual conversion or gametocyte formation. For this purpose, gametocyte formation of PfPPM2-HA-glmS^NF54^ parasites was assessed, which was induced by the addition of serum free media and treatment with N-acetyl glucosamine (NAG) was provided to kill asexual forms (Flammersfeld et al, 2020). While there was no difference in the case of parental NF54 line, a significant reduction in parasite sexual conversion in PfPPM2-HA-glmS^NF54^ parasites was observed upon PfPPM2 depletion (Fig. 3C). These data were consistent with studies performed in *P. berghei*, which implicated PPM2 in gametocyte formation (Guttery et al, 2014). The transcription factor AP2-G is a master regulator of sexual commitment, which is kept repressed by HP1 and H3K9me3 (Filarsky et al, 2018; Josling et al, 2020), and induction of its expression is critical for conversion of asexual to sexual forms (Kafsack et al, 2014). AP2-G expression was determined at the schizont stage by qRT-PCR, which revealed that PfPPM2 depletion significantly reduced AP2-G expression (Fig. 3D) and corroborated well with impaired sexual conversion (Fig. 3C).

HP1 associated H3K9me3 occupies the heterochromatin regions and represses the expression of heterochromatic genes. PfAP2-G is one of the few euchromatic genes that are silenced by HP1-H3K9me3, which is critical for sexual commitment (Brancucci et al, 2014). The impact of PfPPM2 depletion on HP1 and H3K9me3 at the promoter of AP2-G was assessed by ChIP-qPCR at the schizont stage (Brancucci et al, 2014). Results from these assays revealed that PfPPM2 depletion caused a significant increase in both HP1 as well as H3K9me3 occupancy at AP2-G loci (Fig. 3E). Occupancy on gametocyte specific gene Pfs16, which was used as a control and is not regulated by AP2-G or H3K9me3, did not show any difference. Furthermore, Western blot was also performed to check the levels of H3K9me3 in the case of PfPPM2 depletion. A significant increase in H3K9me3 was also observed under PPM2 depletion (Fig. 3F), which is likely to enhance HP1 deposition on the genome and formation of heterochromatin (Bannister et al, 2001). These observations suggested that PfPPM2 may prevent the association of HP1 and H3K9me3 with heterochromatin, which de-represses the expression of AP2-G and promotes sexual differentiation.

Collectively, these observations indicated that the regulation of HP1 by PfPPM2 is important for AP2-G derepression and sexual differentiation of the parasite.

### Optimal HP1-S33 phosphorylation levels are critical for asexual development and sexual differentiation of the parasite

The possibility of PfPPM2 mediated regulation of HP1 playing a role in asexual development of the parasites was also investigated. For this purpose, HP1 was episomally expressed in PfPPM2-HA-glmS parasites (PfPPM2-HA-glmS^Nf54:^HP1-GFP^OE^) (Fig. 4A). Growth rate assays revelaed that HP1 overexpression significantly reversed the defects in parasite growth that were observed upon PfPPM2 depletion (Fig. 4A). In addition, parasite division-indicated by the number of merozoites or nuclear centers per schizonts which is impaired in PfPPM2-HA-glmS^NF54^ parasiteswas also significantly restored (Fig. 4B). Preliminary experiments also indicated that HP1 overexpression reverted the impaired gametocyte formation as a result of PfPPM2-depletion (Fig. S6B). Next, we attempted to mutate the PfPPM2-target site in the endogenous HP1 locus to generate S33 phosphodeficient (S33A) and phosphomimetic mutants (S33D). These efforts yielded a mixed population that contained parasites with S33A mutation along with wild type unmodified HP1 (Supp. Fig. S7) (PfPPM2-3xHA-GlmS^NF54:HP1/HP1-S33A-Flag^), which could be distinguished by IFA with anti-Flag antibody. We were unable to generate S33D mutant parasites which could be due to deleterious effects of this mutation. The flag-tagged HP1-S33A parasites were extremely slow growing and possessed gametocyte-like morphology. IFA revealed that ∼60% of S33A mutant Flag-tagged parasites were gametocytes as they expressed gametocyte specific protein Pfs16. In contrast, most WT HP1-Flag parasites remained asexual (Fig. 4C). These data were consistent with the fact that PfPPM2 mediated dephosphorylation at S33 of HP1 may promote sexual conversion of the parasite.

**Figure 4:**
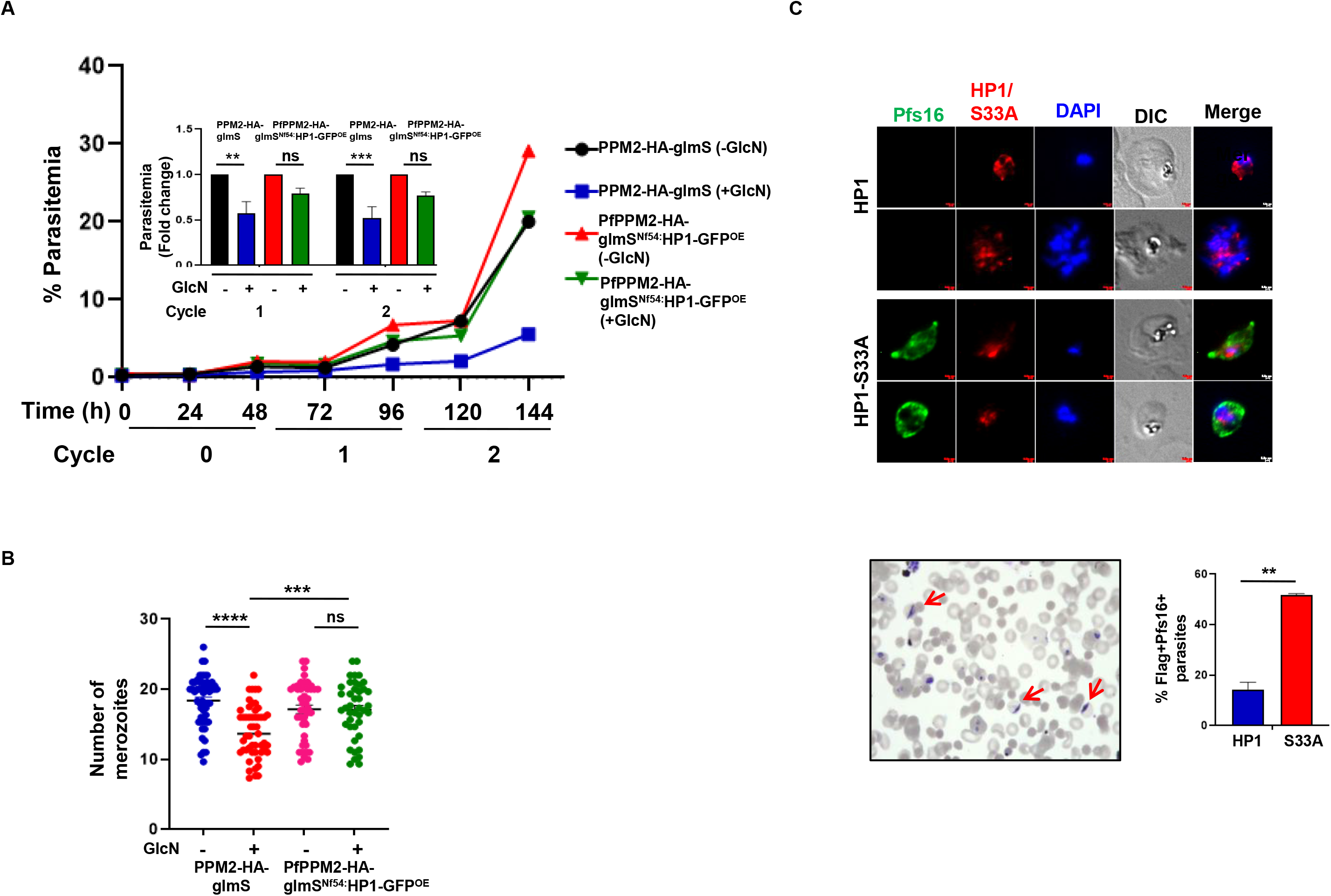
PfPPM2 mediated HP1 phosphorylation may be critical for asexual development and sexual differentiation. A. PfPPM2-HA-GlmS^NF54^ parasites or HP1-GFP overexpressing PfPPM2-HA-glmS^Nf54:^HP1-GFP^OE^ parasites were synchronized, treated with GlcN (cycle 0) and growth rate assays were performed as described for Fig. 1C. *Inset*, fold change in parasitemia was determined with respect to untreated parasites in the next two cycles by performing flow cytometry (SEM ± SE, n=3, ANOVA, **P<0.01, P >0.05, ***P<0.001, ns-non-significant). B. PfPPM2-HA-GlmS^NF54^ or PfPPM2-HA-glmS^Nf54:^HP1-GFP^OE^ parasites were treated with GlcN as described in panel B. Thin blood smears were made ∼40-44 hpi in cycle 1. The number of nuclear centers/merozoites were counted from Giemsa-stained thin blood smears (SEM ± SE, n=3, ANOVA, ****P<0.0001, ***P<0.001, ns-non-significant) C. IFA was performed on PfPPM2-3xHA-GlmS^NF54: HP1/HP1-S33A-Flag^ parasites in which WT-HP1 or its S33A mutant (Supp. Fig. S7) were tagged with Flag. Anti-Flag antibody was used to detect parasites with tagged proteins and anti-Pfs16 was used to stain gametocytes. A representative Giemsa-stained smear of S33A mutant parasites with a significant number of gametocytes (red arrows). *Right Panel,* % Flag-tagged parasites that exhibited Pfs16 expression in IFA (Mean ± SEM, t-test, N=3; **, P<0.01). Scale bar =1 μm.

To evaluate this further, we attempted episomal overexpression of HP1 S33A and S33D mutants in PfPPM2-HA-glmS parasites. Since multiple attempts to either episomally express S33A or S33D mutants were unsuccessful, HP1 phosphomutants tagged with GFP were conditionally overexpressed by fusing with the FKBP Death Domain (DD) at its C-terminus (Supp. Fig. S8A), which can be stabilized by the addition of Shield-1 (Armstrong & Goldberg, 2007). Western blotting revealed successful over-expression of the recombinant proteins (HP1/S33A/S33D-GFP-DD) in the presence of Shld-1 but some amount of proteins was also expressed in its absence and was indicative of "leaky" expression (Fig. S8B). Growth rate assays revealed that both S33A and S33D mutant expressing parasites impaired asexual development as the parasitemia for these parasite lines did not increase and was much lower in cycles 1 and 2 in comparison to WT-HP1 expressing parasites (Fig. 5A). Since these defects were observed in the absence of Shld-1, even low levels of these mutant proteins seemed to impair asexual parasite development.

**Figure 5.**
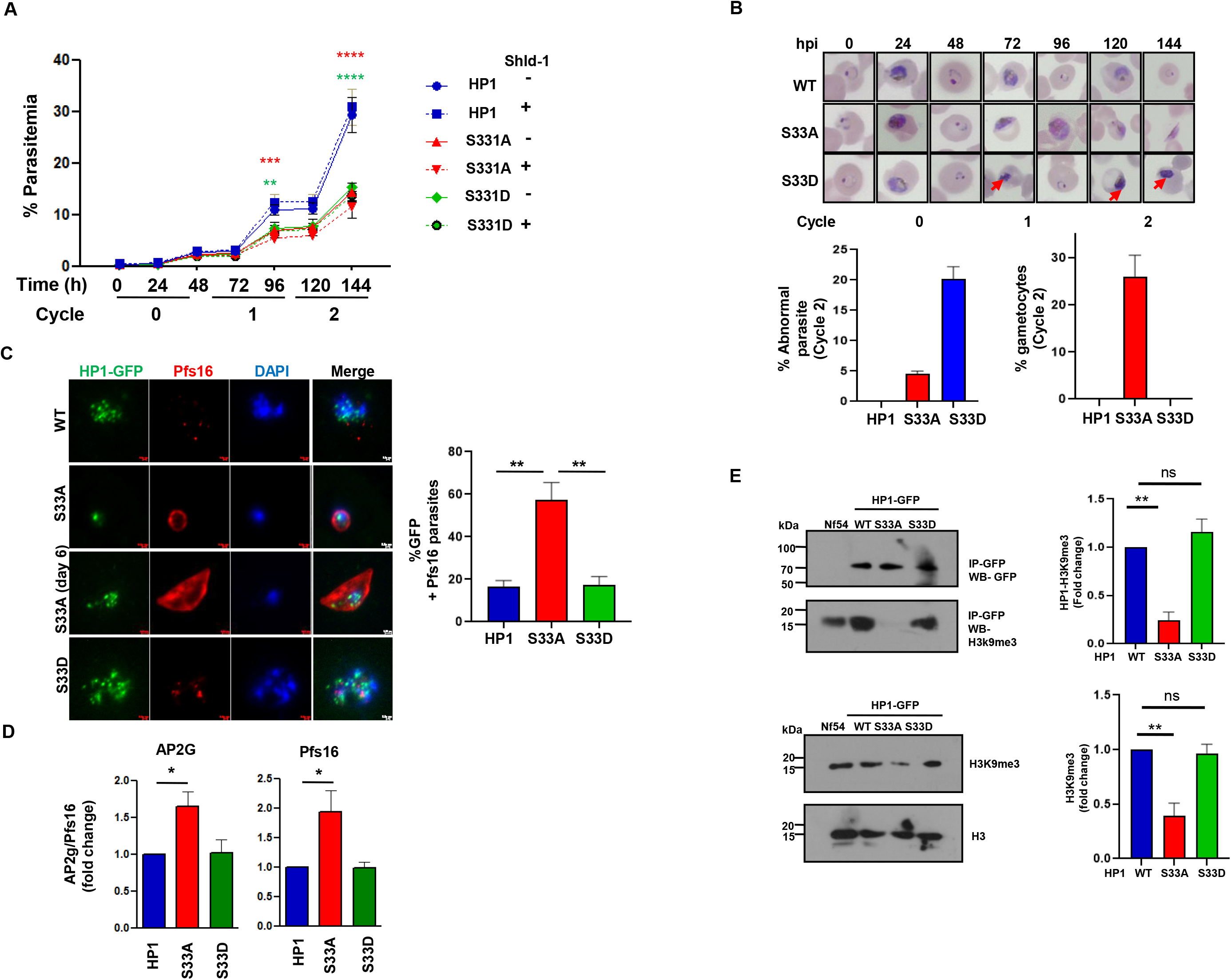
Phosphorylation state of HP1 at S33 is critical for asexual and sexual development of *P. falciparum*. A. HP1/S33A/S33D-GFP-DD parasites-which were generated by overexpressing HP1 or its S33A/D mutants with a DD domain and GFP (Supp. Fig. S8 A,B) -were synchronized and ring stage parasites were used for setting up growth rate assay in the presence or absence of Shld-1. Parasite growth was assessed after each cycle at the indicated time points from Giemsa-stained thin blood smears of parasite cultures and are represented as % of total number of parasites (SEM ± SE, n=3, ANOVA, **P<0.01, ***P<0.001, ****P<0.0001). B. Thin blood smears were made for HP1/S33A/S33D-GFP-DD parasites cultured in the presence of Shld1 at indicated time point and stained with Giemsa to assess their morphology. Gametocytes dominated S33A cultures from cycle 1 and in the case of S33D parasites several parasites exhibited abnormal morphology (red arrows). *Bottom Panel,* % parasites with abnormal or gametocyte morphology was determined. Data from two independent biological replicates is shown. C. IFA was performed on thin blood smears of HP1/S33A/S3D-GFP-DD parasites cultured in the presence of Shld1 using anti-GFP and anti-Pfs16 antibodies. *Right Panel*, % of GFP positive parasites that exhibited Pfs16 staining was determined (SEM ± SE, n=3, ANOVA, **P<0.01). Scale bar = 1 μm. D. qRT-PCR was performed on Shld-1 treated HP1/S33A/S33D-GFP-DD parasites in cycle 1 to assess the fold change in AP2-G and Pfs16 expression with respect to WT-HP1 overexpressing parasites (SEM ± SE, n=3, ANOVA, *P<0.05) E. Nuclear protein lysates prepared from NF54 or HP1/S33A/S33D-GFP-DD parasites cultured in the presence of Shld-1 were used for immunoprecipitation using anti-GFP antibody. The IP (upper panel) or protein lysate (bottom panel) were used for Western blotting using indicated anti-GFP or anti-H3K9me3 and anti-H3 antibodies. *Upper Right Panel*, H3K9me3 co-immunopreciptated with HP1 or S33A/S33D mutants was quantitated by densitometry after normalization with respect to H3K9me3 present in total lysate. Fold change in H3K9me3 present in IP of S33A/S33D mutants with respect to WT HP1 was determined (SEM ± SE, n=3, ANOVA, **P<0.01, ***P<0.001, n=3). *Bottom Right Panel,* Fold change in H3K9me3 levels was determined in S33A/D mutants with respect to WT HP1 in HP1 expressing parasites after normalization with respect to total H3 (SEM ± SE, n=3, ANOVA,**P<0.01, P > 0.05, ns).

The analysis of S33D mutant, which represents S33 hyperphosphorylated form of HP1, overexpressing parasites revealed arrested development as only a small population of trophozoites progressed to schizonts and a signficant number of parasites exhibited abnormal pyknotic morphology (Fig. 5B). These data suggested that S33D mutant causes cell cycle arrest and is possibly toxic to the parasites. In stark contrast, the majority of the S33A-overexpressing parasites had gametocytes like morphology (Fig. 5B), which was confirmed with IFA using antibodies against gametocyte specific protein Pfs16 (Fig. 5C). While only a small fraction of WT-HP1 and S33D overexpressing parasites exhibited Pfs16 staining, in the case of S33A, more than 60% GFP+ cells were Pfs16+. Moreover, these gametocytes developed further to stage IV-V after six days (Fig. 5C). These data confirmed that S33A mutation results in gametocyte formation, which explained slow growth rates of the S33A-overexpressing parasites (Fig. 5A). qRT-PCR revealed that the expression of AP2-G as well as Pfs16 was significantly higher in S33A mutant overxpressing parasites (Fig. 5D), which corroborated well with the fact that these parasites were mainly gametocytes. Collectively, these data suggested that dephosphorylation of HP1 at S33 by PfPPM2 activation-which is mimicked by the S33A mutant- is critical for sexual differentiation of the parasite as PPM2 depletion prevents this process. While phosphorylation of HP1-S33 is critical for parasite asexual divsion, its hyperphosphorylation mimicked by S33D mutant-(which to some extent represents PfPPM2-depleted state)can cause arrest in parasite development.

Next, using the S33A/D mutants, efforts were made to investigate if S33 dephosphorylation regulates HP1-H3K9me3 interaction, which is needed for maintenance of heterochromatin at AP2-G and other locus (Brancucci et al, 2014). To this end, immunoprecipitation was performed to assess the interaction between WT or mutant HP1 with H3K9me3 using these transgenic parasites. GFP-HP1 was able to pull down H3K9me3, as reported previously (Flueck et al, 2009). Strikingly, there was a dramatic loss of its interaction upon S33A mutation (Fig. 5E). In contrast, the amount of H3K9me3 pulled down with S33D mutant was almost unaltered. Furthermore, Western blotting of nuclear lysate indicated that levels of H3K9me3 were also reduced in the case of S33A mutant in comparison to WT or S33D overexpressing parasites (Fig. 5E). These data suggested that dephosphorylation of S33 prevents the interaction of HP1 with H3K9me3 and may also impair trimethylation of H3K9, which results in the expression of AP2-G and sexual differentiation of the parasite. Therefore, dephosphorylation of pS33-HP1 is a signal for sexual commitment. Consistent with this, IFA revealed that most WT HP1 or S33D overexpressing asexual parasites were stained with anti-pS33-HP1. In contrast, pS33-HP1 was absent from almost all S33A-overexpressing parasites, which were mainly gametocytes (Fig. 6A). Next, the effect of induction of sexual conversion by serum removal on pS33-HP1 levels was assessed (Fig. 6B). IFA revealed that more than half of the PfPPM2-HA-glmS^NF54^ schizonts exhibited pS33-HP1 staining but upon induction of sexual conversion most of the parasites that were either gametocytes and almost all schizonts-possibly sexually committed forms-lost pS33 phosphorylation (Fig. 6B). In contrast, upon PfPPM2 depletion, mostly shizonts were observed even after induction that exhibited pS33 staining. Given that PfPPM2-depletion prevents sexual differentiation, it is reasonable to suggest that these pS33-HP1-positive schizonts are unable to undergo gametocytogenesis due to lack of PfPPM2-mediated dephosphorylation of this site. Furthermore, in an unrelated parasite line, pS33-HP1 staining was also lost upon conversion to gametocytes and was observed mainly in asexual parasites (Supp. Fig.S10). Collectively, these data suggested that dephosphorylation of S33 of HP1 by PfPPM2 is a signal for sexual differentaition of the parasite, which triggers epigenetic changes necessary for this process (Fig. 6C).

**Figure 6:**
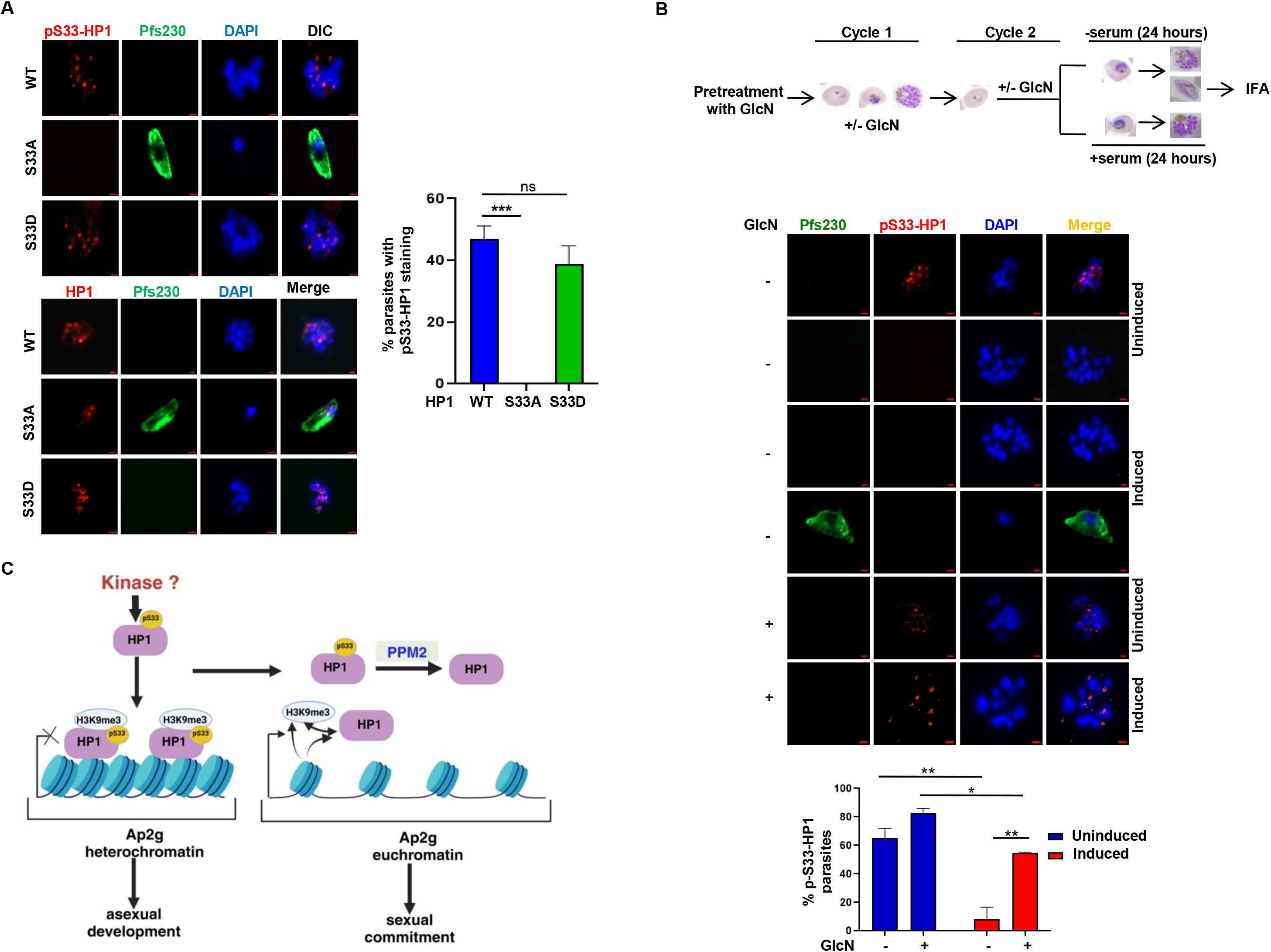
Dephosphorylation of HP1 is a novel signal for sexual differentiation of the parasite. A. IFA was performed on thin blood smears of HP1/S33A/S33D-GFP-DD parasites cultured in the presence of Shld-1 using anti-pS33-HP1 and anti-Pfs230 antibodies, revealed that pS33-HP1 was present mainly in the asexual stages (HP1/HP1S33D) but it was absent from HP1-S33A parasites, majority of which were gametocytes (SEM ± SE, n=3, ANOVA, ***P<0.001, ns-non-significant). Scale bar = 1 μm. B. Sexual conversion of PfPPM2-HA-glmS^NF54^ cultured in the absence or presence of GlcN was induced by serum depletion as indicated in the schematic. IFA was performed on parasites pre- and post-induction using anti-pS33-HP1 and anti-Pfs230 antibodies. % pS33-HP1 positive parasites in each condition were determined (SEM ± SE, n=2, ANOVA, *P<0.05,**P<0.01, P > 0.05, ns). Scale bar = 1 μm. C. Schematic illustrates a novel signalling pathway in *P. falcipraum*. Present studies demonstrate that PfPPM2 is important for both asexual division and sexual differentiation of *P. falciparum*. A yet to be identified kinase phosphorylates HP1 at S33 which is important for asexual development and division. S33-HP1 phosphorylation promotes its association with H3K9me3 and maintenance of heterochromatin state critical for asexual division. PfPPM2 mediated dephosphorylation of HP1 at S33 prevents interaction of HP1 with H3K9me3 and promotes euchromatin formation and derepression of AP2-G, which is critical for sexual differentiation.

## Discussion

Present studies demonstrate that protein phosphatase PfPPM2 is a versatile enzyme as it regulates both asexual and sexual development of *P. falciparum*. It is interesting to note that its *P. berghei* homologue is dispensable for asexual development, which suggested that PPM2 may regulate diverse processes in these two *Plasmodium* spp, alternatively, other PPs may complement its function in *P. berghei*. While PbPPM2 disruption results in reduced gametocytemia as well as lower female:male gametocyte ratio as well as ookinete differentiation (Guttery et al, 2014), however, the mechanism via which PPM2 regulates parasite development had remained unclear. Conditional depletion of PfPPM2 revealed that it regulates the division of the *P. falciparum* and the phosphoproteomics studies revealed that PfPPM2 influence the phosphorylation of several proteins implicated in epigenetic regulation and translation (Table S1), which included histone 3 and 3.3 and HP1. We went on to confirm that the dephosphorylation of histone H3 at S10 and S28 and that of HP1 at S33 is regulated by PfPPM2. The phosphorylation of histone H3 N-terminal tail at S10 and S28 in mammals and yeast has been associated with mitosis and transcriptional control, respectively (Birnbaum et al, 2017; Salcedo-Amaya et al, 2009; Sawicka et al, 2014; Sawicka & Seiser, 2012). Interestingly, H3 phosphorylation regulates distinct processes like stabilizing chromosome condensation during mitosis which contributes to transcriptional repression and activation during the interphase (Perez-Cadahia et al, 2009). In addition, these sites need to be dephosphorylated by phosphatases like PP1 in mammalian cells failing which there is mitotic arrest (Bosch et al; Gil & Vagnarelli, 2019; Qian et al, 2011). While phosphorylation of these sites has been reported in *P. falciparum* (Dastidar et al, 2013), there is almost no information on their role in the parasite biology. It is possible that dephosphorylation of Histone H3 and possibly its variant H3.3 by PfPPM2 may also be critical for parasite division. Interestingly, transcriptome analysis revealed that several genes implicated in cell division like Centrin-1, Centrin-4, PCNA and genes involved in translation were deregulated upon PfPPM2 depletion, which was consistent with observed defects in PfPPM2-depleted parasites (Supp. Fig. S9), which hinted at epigenetic changes as a result of aberrant phosphorylation of above-mentioned proteins. The fact that HP1-overexpression prevented defects in asexual development and parasite division (Fig. 4A,B) also supports that chromatin remodelling via phosphorylation of proteins like HP1 and histone H3 contributes to parasite division. We observed defects in MTOC/CP biogenesis in PfPPM2-depleted parasites (Fig. 1F,G). Since no major targets of PfPPM2 identified by phosphoproteomics relate to these processes, it is reasonable to suggest these are likely to be an outcome of arrested division.

Strikingly, H3K9me3 levels were significantly higher in PfPPM2-depleted parasites (Fig. 3F). In *P. falciparum*, H3K9me3-marked heterochromatin along with HP1 contributes to the repression of genes like AP2-G, which is involved in sexual commitment and selective expression of var gene family members that code for proteins exported to the host surface and are involved in evasion from the host immune system. AP2-G, which is involved in sexual commitment, is one of the few euchromatic genes which is silenced by HP1(Brancucci et al, 2014; Kafsack et al, 2014). Consistent with higher H3K9me3 (Fig.3F), the expression of AP2-G was reduced upon PfPPM2 depletion (Fig.3D), which could be explained by enhanced occupancy of both H3K9me3 and HP1 on AP2-G locus (Fig. 3E). PfHP1-H3K9me3 interaction is critical for maintaining the H3K9me3 marks, which represses the expression of genes like AP2-G and prevents sexual differentiation. HP1 also regulates mutually exclusive expression of Var genes present at heterochromatic loci and in turn contributes to antigenic variation of PfEMP1 (Brancucci et al, 2014). Since present studies were focussed on asexual and sexual development of the parasite, regulation of var genes was not investigated, although transcriptome analysis suggested several var genes that normally exhibit mutually exclusive expression and were aberrantly expressed upon PfPPM2 depletion (Supp. Fig. S9).

The PfPPM2 target phosphorylation site S33 on HP1 is present in its chromodomain via which it interacts with H3K9me3 (Bannister et al, 2001; Bui et al, 2021). Phosphodeficient (S33A) and phospho-mimic (S33D) mutants-that resemble high or low PfPPM2 activity-were used to demonstrate that dephosphorylation of S33 is important for HP1-H3K9me3 interaction. PfPPM2 mediated dephosphorylation-mimicked by S33A mutant-may disrupt this interaction resulting in higher AP2-G expression (Fig. 5D,E), which is critical for sexual commitment of the parasite as it facilitates the expression of AP2-G and downstream genes needed for this purpose (Kafsack et al, 2014; Llora-Batlle et al, 2020). It was also interesting to note that H3K9me3 levels were much lower in S33A mutant (Fig.5E), which was consistent with higher H3K9me3 in PfPPM2-depleted parasites (Fig. 3F). Previous studies have indicated that upon binding H3K9me3, HP1 also recruits histone methyl transferases (HKMTs) to enhance H3K9me3 levels (Loyola et al, 2009).

These data explained the reduced expression of AP2-G in PfPPM2-depleted parasites (Fig. 3D) that exhibit impaired sexual conversion (Fig. 3C), which was supported by the fact that S33A mutant overexpressing parasites-resembling PfPPM2 activated state-constitutively expressed AP2-G (Fig. 5D) and promoted sexual conversion (Fig. 5B,C). Consistent with this notion, pS33-HP1 was mainly detected in asexual parasites and was absent from gametocytes (Fig. 6A,B). Importantly, PfPPM2-depletion caused a significant increase in pS33-HP1 in asexual parasites (Fig. 6B) and corroborated well with reduced gametocytes formed by these parasites (Fig. 3C). Therefore, dephosphorylation of HP1 was identified as a novel signal which promotes sexual differentiation of malaria parasite by above-mentioned mechanism (Fig. 6C). It will be important to identify the kinase which phosphorylates HP1-S33 to promote asexual division and prevent sexual conversion. Given that HP1 overexpression overcomes defects in parasite division as a result of PfPPM2-depletion (Fig. 4A and 4B) suggests that the regulation of HP1 may also be critical for parasite division. Previous studies indicate that HP1 depletion also results in parasite cell cycle arrest (Brancucci et al, 2014). Interestingly, HP1-S33D mutant, which mimics hyperphosphorylated form of HP1 at S33, also exhibited severe growth arrest. Therefore, PfPPM2 keeps phosphorylation of S33 to optimal levels, which seem to be necessary for asexual development of the parasite-its hypophosphorylation results in sexual differentiation whereas hyperphosphorylation causes arrest in asexual development (Fig. 6C).

Present studies also indicate that PfPPM2 promotes mRNA translation or protein synthesis by preventing eIF2α phosphorylation by keeping eIF2α kinase PK4 in check (Supp. Fig. S5). Alternatively, it may directly dephosphorylate eIF2α.Given that protein synthesis is important for cell cycle progression (Polymenis & Aramayo, 2015), it is possible that PfPPM2 may regulate parasite division by regulating this process.

## Experimental Procedures

Information related to reagents and detailed methods are provided in Experimental Procedures provided in the Supplementary Information.

### Parasite cultures and Transfection

*Plasmodium falciparum* 3D7 and NF54 parasites were cultured in O^+^ human erythrocytes in RPMI-1640 supplemented with 0.5% AlbumaxII and 50μg/mL hypoxanthine as described previously (Trager & Jensen, 1977) (Lambros & Vanderberg, 1979). For transfection, ring stage parasites were electroporated with ∼100 μg plasmid DNA constructs (details provided in Supplementary Experimental Procedures). Parasites were maintained initially on 2-10nM WR99210 followed by 400 μg/ml of G418.

### Assays growth rate, parasite division and gametocyte formation

*PfPPM2-HA-glmS* parasites were cultured in the presence of 10nM WR99210 and 400μg/ml G418 and *PfPPM2-HA-glmS^Nf54:^HP1-GFP*^OE^ parasites were cultured in the presence of 10 nM WR99210, 400μg/ml G418 and 2.5μg/ml of Blasticidin. For the depletion of PfPPM2, ring stage parasites were typically treated with 2.5mM glucosamine (GlcN) for one cycle and parasite growth was assessed at an interval of 24-h for additional 144-h. For determining parasitemia, using flow cytometry samples were fixed with 1% PFA and 0.0075% glutaraldehyde solution samples were either stored at 4°C or processed directly for Hoechst 33342 staining for 10-min at 37°C and analysis was performed on a BDverse (BD biosciences) (Rawat et al, 2023; Theron et al, 2010). The data were processed and using FlowJo software. Growth of the parasites was also determined by counting parasites from Giemsa-stained thin blood smears of the cultures. For assessing parasite division, thin blood smears of mature schizont parasites were prepared, microscopically examined, images were captured. The numbers of merozoites from at least 50 schizonts for each condition, per replicate, were counted and average number of merozoites per schizont was determined.

For inducing gametocytogenesis, sexual conversion was induced as previously described with some minor modifications (Brancucci et al, 2017; Flammersfeld et al, 2020). Parasites were seeded at 2% ring parasitemia in 5% haematocrit. In the next cycle, when parasitemia reached ∼10%, gametocytogenesis was induced by replacing serum containing complete RPMI1640 (cRPMI) medium with serum free medium. After ∼24h, serum free medium was replaced by cRPMI treatment with 50mM N-acetyl-D-Glucosamine, which was provided for 5 days to eliminate the asexual parasites unless indicated otherwise. Giemsa smears were made periodically and the number of gametocytes was counted at day 5 p.g.i.

### Immunofluorescence Assays (IFAs)

Immunofluorescence assays (IFA) were performed on thin blood smears as previously described (Tonkin et al, 2004). Briefly, air-dried thin blood smears were fixed with cold methanol and acetone mix (1:1, v:v) for 2-min followed by blocking with 3% BSA for 45-min at room temperature prior to incubation first with primary antibodies and then with secondary antibodies. Finally, Vecta shield mounting media was used (Vector Laboratories Inc.), which contained DAPI to label nuclei. Fluorescence microscopy was performed using Axio Imager Z1 microscope or a LSM980 confocal microscope (Carl Zeiss).

### Immunoprecipitation and Immunoblotting

Immunoprecipitation was performed on nuclear lysates. For making the nuclear lysate, fractionation was performed as previously described (Filarsky et al, 2018; Flueck et al, 2009) with some minor modifications. 200 µg of protein lysate (nuclear extracts) was typically used and incubated with relevant antibodies at 4°C for 12-h followed by incubation with protein A+G sepharose beads for 5-h. After 3 washes with wash buffer, beads were boiled at 95°C for 10-min. After boiling, supernatant was used for immunoblotting along with the input. After electrophoresis on SDS-PAGE, lysate proteins were transferred to a nitrocellulose membrane and immunoblotting was performed as described previously (Ekka et al, 2020; Kumar et al, 2017).

### qRT-PCR

RNA isolation was carried out using TRIzol reagent (G Biosciences). Real time PCR was carried out using CFX96 Real Time PCR detection system (Biorad). For qRT-PCR, 1 μg of DNAse free RNA was used for cDNA synthesis using Verso cDNA synthesis kit (Thermo scientific), as per the manufacturer’s recommendation. Random primers supplied with the kit were used for the cDNA synthesis. 18srRNA was used as a housekeeping gene for normalization.

### ChIP-qPCR

PfPPM2-HA-glmS^NF54^ parasites were cultured in the presence (2.5 mM GlcN) or absence of glucosamine (GlcN) and were harvested at the end of first cycle at ∼90 h.p.i by fixing the cells. Isolation of chromatin after fixation with formaldehyde was done followed by immunoprecipitation using anti-HP1 or anti-H3K9me3 antibodies or rabbit IgG which was used as negative control as described previously (Brancucci et al, 2014). ChIP or input DNA was used for qPCR (see Supplementary Information for detailed Experimental Procedures).

### Phosphoproteomics and Transcriptomics

The 3D7 or PfPPM2-HA-glmS^3D7/Nf54^ parasites were cultured and synchronized using sorbitol. Ring stage parasites were treated with glucosamine for 72h. The parasites were subsequently harvested upon maturation to schizonts in cycle 1 (∼44 hours post-infection), by lysing the infected red blood cells with saponin. Subsequently, parasite pellets were used independently for RNAseq (PfPPM2-HA-glmS^Nf54^) or phosphoproteomic (PfPPM2-HA-glmS^3D7^) studies. A detailed account of these studies is provided in Supplementary Experimental methods.

## Supporting information

Suuplementary Methods, Tables S1-S3, Figures S1-S10

## Acknowledgements

Studies were supported by grant CRG/2021/000147 from Science and Engineering Research Board (SERB), Department of Science and Technology, India and a Team Science grant (IA/TSG/21/1/600261) to PS and TSKP from DBT-Wellcome Trust India Alliance and funds from NII core. P.S. is a recipient of J.C. Bose Fellowship. The efforts made by Devanshi Sharma for her assistance in RNAseq data analysis and Manish Kumar in critically reading the manuscript are appreciated.

## Supplementary Material

Materials and Methods,

Supplementary Figures S1-S10,

Supplementary Tables S1-S3

(Single PDF)

Datasets S1-S3

(3 Excel files).

## Notes

### Competing Interest Statement

The authors have declared no competing interest.

