## Supplementary material for "PfPPM2 signalling regulates asexual division and sexual conversion of human malaria parasite *Plasmodium falciparum*": Suuplementary Methods, Tables S1-S3, Figures S1-S10

### Material and Methods

*Reagents and antibodies*

Fine chemicals were purchased from Sigma-Aldrich®/MercK including Glucosamine hydrochloride (Cat. No.- G1514), E64 protease inhibitor (Cat. No.- E3132), N-acetyl-D-glucosamine (Cat. No,- A8625), Puromycin (Cat. No.-P8833), Fatty acid free BSA (Cat. No.- A6003-), Oleic acid (Cat. No,- O1008), Palmitic acid (Cat. No.-P0500), Blasticidin (Cat. No.-15205) and G-418-Disulphate (Geneticin Sulphate)(Cat. No.-3341)(TM Media). Oligonucleotides were procured form Sigma-Aldrich®, restriction enzymes were obtained from New England Biolabs, USA. Commercially available antibodies were obtained from Santa Cruz Biotechnology® mouse antiβ-Actin (C4, sc-47778 HRP), anti-Flag (Cat. No.- sc-51590) and Roche mouse anti-GFP (SKU-11814460001) and Roche mouse anti-HA (SKU-11583816001), anti-centrin (20H5 monoclonal mice antibody, Cat. No.-04-1624), anti-α-tubulin antibody (mouse monoclonal, clone DM1A, Cat. No.- T6199), anti-Puromycin (Cat. No.- MABE343). Antibodies were also procured from, Cell Signaling Technology, Inc (CST)- Tri-Methyl-Histone H3 (H3K9me3) (D4W1U) Rabbit (Cell signalling and technology), Histone H3 (D2B12) XP® Rabbit mAb (ChIP Formulated) (4620S), eIF2A-D7D3 rabbit mAb, CST, from abcam- anti-H3S10ph (ab5176), anti-H3S28ph (ab5169) and Thermo scientific- Phospho-EIF2S1 (Ser51) Antibody (PA5-37800). Protein A/G plus agarose beads (sc-2003) were obtained from Santa Cruz Biotechnology®. Shield-1 was purchased from Cheminpharma, LLC, USA. Anti-PfMSP1 (MRA-34), Anti-Pfs16 (MRA-1276), Pfs230 (MRA-878A) and DSM1 were obtained from *BEI* resources, Malaria Research and Reference Reagents Resource Centre (MR4). pS33-HP1 and non-phospho-HP1 was custom synthesized from Antagene.

Proteomics related reagents: Sigma Aldrich - Triethylammonium biocarbonate (TEAB)(Cat. No.-T7408), Dithiothreitol (DTT) (Cat. No.- D9779), Iodoacetamide (IAA) (Cat. No.- I1149), Pierce™ - BCA estimation kit (Cat. No.- 23225), Quantitative Peptide Assays & Standards (Cat. No.- 23275), TMT10 plex Labelling Reagent (1 X 0.8 mg) (Cat. No. 90110), Trypsin/Lys-C mix Promega, Mass spec grade (Cat. No.- V5073), Waters SepPak cartridges Vac 1cc 50mg (Cat. No.- WAT054955), Water for chromatography LC-MS (Cat. No.- 1.00029.4000), Acetonitrile hypergrade for LC-MS (1.15333.4000), Formic acid (Cat. No.- 94318), High Select™TiO2 Phosphopeptide Enrichment Kit (Cat. No.- A32993), High Select™Fe-NTA Phosphopeptide enrichment kit (Cat. No.- A32992),Hydroxylamine (Cat. No.- 90115).

**Parasite cultures**

*Plasmodium falciparum* strains 3D7 and NF54 were obtained from *BEI* resources, Malaria Research and Reference Reagents Resource Centre (MR4), American Type Culture Collection (ATCC). Parasites were maintained in $O^+$ human erythrocytes (5% hematocrit) in RPMI-1640 supplemented with 0.5% AlbumaxII and 50µg/mL hypoxanthine (cRPMI1640). Cultures were maintained at 37°C under a mixture of 5% $CO_2$, 3% $O_2$, and 91.8% $N_2$ or 5% $CO_2$. Fresh erythrocytes were used to dilute parasites with fresh culture media to maintain 3-5% parasitemia at 5% haematocrit (Trager & Jensen, 1977). For culturing various transgenic lines, relevant drugs were used as described above. Parasite synchronization was carried out using sorbitol as described previously (Lambros & Vanderberg, 1979).

**Generation of plasmid constructs and transfection of parasites**

*PfPPM2-3xHA-GlmS*$^{3D7/NF54}$

In order to fuse *PfPPM2* with *glmS* and introduce a 3xHA tag at the C-terminus, a homology region corresponding to its 3'-end PfPPM2 was PCR amplified using primers (1/2) from *P. falciparum* 3D7 genomic DNA and cloned in pGlmS-SLI vector (a kind gift from Prof. Alan Cowman and further modified by Dr. Rahul Rawat by incorporating 2A skip-NeoR cassette) using sites for *BglII* and *PstI.* Subsequently, PfPPM2-3xHA-glmS construct was then transfected in 3D7 and NF54 parasites. For the transfection of this and other plasmids described below, 8–10% ring stage parasites were electroporated with ~100 μg plasmid DNA. Parasites were maintained initially on 2-10 nM WR99210 followed by 400 μg/ml of G418 for selection linked integration. A mixed population of parasites, which included both integrants and wild type parasites, was observed upon genotyping of drug-selected parasites. The mixed population was subjected to limited dilution cloning, the clones obtained after cloning were verified for correct 5' and 3' integration and the absence of the wild type unaltered region by genotyping PCR. PCR amplicons were sequenced for further confirmation. Genotyping was performed by using sets of PCR primers indicated in Supplementary Table S2.

*PfPPM2-3xHA-GlmS*$^{NF54:\ HP1/HP1\text{-}S33A\text{-}Flag}$

To generate pSLI-3x-Flag-yDHODH-BSD vector, first amplification of 2A-yDHODH fragment was done from pSLI-N-sandwich loxP vector (Birnbaum et al, 2017) using primer (7/8) and cloned using *KpnI* and *XhoI* sites in pSLI-HA-glmS-BSD (a kind gift from Dr. Asif Mohmmed lab and further modified by Dr. Rahul Rawat by incorporating 2A skip-NeoR cassette) vector to yield pSLI-3x-Flag-2A-yDHODH-BSD, which resulted in the replacement of the 3xHA-tag by 3xFlag coding sequence and introduction of 2A skip peptide along with yDHODH cassette and removal of NeoR and glmS sequence. For tagging HP1 at the C-terminal end with 3x Flag tag, 3'- homology region was amplified from 3D7 genomic DNA using primer (9/10) and cloned using *SpeI* and *KpnI* sites to generate pSLI-HP1-3x-Flag-yDHODH-BSD construct. For generating, S33A-mutant tagged with flag, pcr primers (11/10)-in which codons for S33 were replaced with those for A-were used to clone the homology region using *SpeI* and *KpnI* sites of pSLI-3x-Flag-yDHODH-BSD vector. This construct was transfected in PPM2-HA-glmS$^{NF54}$ parasite line and subsequently transgenic parasites were selected by using blasticidin (2.5µg/ml) and recombinants were enriched using 1.5µM DSM-1 and drug resistant parasites were genotyped for 5'-and 3'-integration and to check for the unmodified wild type locus of HP1. The desired mutation was confirmed by Sanger sequencing.

*PfPPM2-HA-glmS$^{Nf54:}$HP1-GFP$^{OE}$ or HP1/S33A/S33D-GFP-DD*

HP1 was cloned in either pARL-GFP with BSD (a kind gift from Dr. Mohmmed Asif) or pARL-GFP-DD vector, which was modified as described below. For generating the pARL-HP1-GFP-DD, HP1 was amplified by PCR from 3D7 genomic DNA using primers (16/17) and GFP coding sequence was amplified from cpARL-GFP-BSD vector using primers (18/19) and the destabilization domain (DD) was amplified using primers (20/21) from pTEX-HA-DD vector. The HP1 and GFP domain was fused by overlapping PCR using primer (16/19) and GFP and DD domain was fused together using primer (18/21). The final overlapping PCR to obtain HP1-GFP-DD fused together was done by using primer set

(16/21) and cloning was performed using *KpnI* and *AvrII* restriction sites of cpARL-GFP-BSD vector. S33A mutations in HP1 were generated by site directed mutagenesis using primers (22/23) and S33D mutation was generated using primers (24/25), which were confirmed by sequencing. Subsequently, mutants were cloned in pARL-GFP-DD vector like the WT HP1. For pARL-HP1-GFP construct, HP1 was cloned in pARL-GFP-BSD vector using primers 16/17' using *KpnI* and *AvrII* sites.

These plasmid constructs were transfected in PPM2-HA-glmS$^{NF54}$ for generating PfPPM2-HA-glmS$^{Nf54:}$HP1-GFP$^{OE}$ or NF54 parasites for HP1/S33A/S33D-GFP-DD parasites and were selected with 2.5μg/ml of Blasticidin.

**Growth rate assays**

*PfPPM2-HA-glmS* parasites were cultured in the presence of 10 nM WR99210 and 400μg/ml G418 and PfPPM2-HA-glmS$^{Nf54:}$HP1-GFP$^{OE}$ parasites were cultured in the presence of 10 nM WR99210, 400μg/ml G418 and 2.5μg/ml of Blasticidin. Subsequently, ring stage parasites were seeded at ~0.5% parasitemia at 2% hematocrit after synchronisation. For the depletion of PfPPM2, ring stage parasites were typically treated with 2.5mM glucosamine (GlcN) for one cycle and parasite growth was assessed at an interval of 24-h for additional 144-h. For growth rate assays with HP1/S33A/S33D-GFP-DD lines, parasites were synchronized and seeded at ~0.5% ring parasitemia and parasites were cultured in the presence or absence of Shield-1 (250nM) for two cycles. Typically, samples were collected after every 24h for making thin blood smears as well as FACS analysis.

For determining parasitemia using flow cytometry, samples were fixed with 1% PFA and 0.0075% glutaraldehyde solution and kept on an end-to-end rocker for 15min. After completion, samples were either stored at 4$^{o}$C or processed directly for Hoechst 33342 staining for 10-min at 37ºC. Samples were then washed at least twice with FACS buffer followed by analysis on BDverse (BD biosciences) for 10,000-100,000 events per sample (Theron et al, 2010). The data was processed and analyzed in FlowJo software.

**Assessment of parasite division**

Parasite lines were synchronized and GlcN treatment was provided for one cycle, E64 (10µM) was added at ~40h h.p.i. (cycle 1) schizonts for 4-5 hours. Thin blood smears were prepared, stained with Giemsa, images of mature schizont parasite were captured on a Leica bright field microscope. The numbers of merozoites from at least 50 schizonts for each condition, per replicate, were counted and average number of merozoites per schizont was determined.

**Gametocyte conversion assay**

For inducing gametocytogenesis, sexual conversion was induced as previously described with some minor modifications (Brancucci et al, 2017; Flammersfeld et al, 2020). Briefly, PPM2-HA-glmS$^{NF54}$ parasites were treated with 2.5 mM GlcN for one cycle (pre-tretament). Subsequently, parasites were seeded at 2% rings in 5% haematocrit. In the next cycle, when parasitemia reached ~10%, gametocytogenesis was induced by replacing serum containing complete RPMI1640 (cRPMI) medium with serum free medium. After ~24-h, serum free medium was removed followed by supplementation of cRPMI and treatment with 50mM N-acetyl-D-Glucosamine was provided for 5 days to eliminate the asexual parasites unless indicated otherwise. Giemsa smears were made periodically and the number of gametocytes was counted at day 5 p.g.i. Sexual conversion rates were determined by counting gametocytes formed at day 5 relative to the initial parasites, which were mainly asexual rings (Coleman et al, 2014; Llora-Batlle et al, 2020). In order to assess the sexual conversion of the HP1/S33A/S33D-GFP-DD parasites, sorbitol synchronization was performed at the ring stage and parasites were left untreated or treated with Shld-1 (250 nM). Subsequently, IFAs were performed after 4-6 days on blood smears using anti-GFP and anti-Pfs16 or anti-Pfs230 antibodies. The number of Pfs16/Pfs230-stained parasites that represented gametocytes was counted. Since pS33-HP1 antibody was raised in rabbits, co-staining was done using anti-Pfs230 which was raised in mice.

**Immunofluorescence Assay (IFAs)**

Immunofluorescence assays (IFA) were performed on thin blood smears as previously described (Tonkin et al, 2004). Briefly, air-dried thin blood smears were fixed with cold methanol and acetone mix (1:1, v:v) for 2-min followed by blocking with 3% BSA for 45-min at room temperature. Subsequently, smears were incubated with primary antibodies for 12-h at 4°C. After washing with blocking buffer, Alexa fluor mouse/rabbit 488/594-labeled secondary antibodies were added (Invitrogen) for 2-h at ambient temperature. Finally, Vecta shield mounting media was used (Vector Laboratories Inc.), which contained DAPI to label nuclei. Fluorescence microscopy was performed using Axio Imager Z1 microscope or a LSM980 confocal microscope (Carl Zeiss). The images were processed using AxioVision 4.8.2 or Zeiss ZEN black/blue software and unless indicated otherwise best representative z-stacks were used for illustrations in the figures.

**Immunoblotting**

Parasite cultures were collected and iRBCs were lysed using 0.05% saponin (w/v) followed by incubation on ice for 10-min. Centrifugation at 8000 rpm was done to isolate the parasite pellet, which was washed three times with pre-chilled PBS. The parasite pellet was re-suspended in lysis buffer (10 mM Tris pH 7.5, 100 mM NaCl, 5 mM EDTA, 1% Triton X-100, and complete protease inhibitor cocktail; Roche Applied Science) or 2% SDS and then homogenized by either passing the solution through a 26-gauge needle or by using sonication at 60% amplitude for 20-sec (1-sec ON/OFF). The supernatant from the centrifugation of the lysates at 14,000 g for 30-min at 4°C was used to estimate the amount of protein using a BCA protein estimation kit (Pierce). Nuclear protein lysates were prepared as described below.

After SDS-PAGE, proteins were transferred to nitrocellulose membrane and membranes with transferred proteins were blocked with 3% BSA/0.2% Tween 20 in 1xTBS (v/v) for 1-h. Primary antibody solutions were made in 3% BSA, diluted to the required concentration and membranes were incubated for 12-h at 4°C. The membrane was then rinsed three times with 1xTBS/0.1% Tween 20 (TBS-T) and incubated with Horseradish Peroxidase (HRP) conjugated secondary antibody prepared in 3% skimmed milk/1xTBS-T or 3%BSA/1x TBS-T (used for phospho antibody-blots) for 2-h. The nitrocellulose membrane was once again washed with 1xTBS-T and detection was performed using West Pico or Femto chemiluminescence substrate from Pierce (USA) after exposure of the membrane to X-ray films.

### Immunoprecipitation

Immunoprecipitation was performed on nuclear lysates. For making the nuclear lysate, fractionation was performed as previously described (Filarsky et al, 2018; Flueck et al, 2009) with some minor modifications. Saponin-lysed infected red blood cells were lysed in ice-cold cell lysis buffer (20mM HEPES pH 7.9, 10mM KCl, 1mM EDTA, 1mM EGTA, 1mM DTT, protease inhibitors) for 5-min. After a 5-min centrifugation at 5200 rpm (2500xg), the cytosolic extract was collected and stored at -80ºC and the nuclei obtained were treated with DNase I and RNAse A [(20mM Hepes pH 7.4, 10mM NaCl, 5mM MgCl2, 1mM CaCl2, 0.1% NP-40, 1x protease inhibitor (Roche Diagnostics), 1x PhosSTOP (Roche Diagnostics), 0.5U/µl DNase I, 1µg RNase A] and incubated in 250 µl reaction volume for 5-min at 37ºC.). Reactions were adjusted to 500 µl using 2x high salt extraction buffer (20 mM Hepes pH 7.4, 1M NaCl, 6mM EDTA, 2mM EGTA, 2mM TCEP, 1x protease inhibitor (Roche Diagnostics), 1x PhosSTOP (Roche Diagnostics), 50% Glycerol, 0.1%NP-40) and nuclear proteins were extracted in 2.5 pellet volume of high salt extraction buffer (20mM Hepes pH 7.4, 500mM NaCl, 3mM EDTA, 1mM EGTA, 1mM TCEP, 1x protease inhibitor (Roche Diagnostics), 1x PhosSTOP (Roche Diagnostics), 25% Glycerol, 0.1%NP-40) by vortexing for 30-min at 4˚C. After centrifugation at 20800g at 4°C for 30-min, supernatant was separated immediately and used for IP experiments after dilution. High salt nuclear extracts were diluted to 250mM NaCl using dilution buffer (20mM Hepes pH 7.4, 1mM EDTA, 1mM TCEP, 1x protease inhibitor (Roche Diagnostics), 1x PhosSTOP (Roche Diagnostics), 25% Glycerol).

For immunoprecipitation, 200 µg of nuclear protein lysate was typically used and incubated with relevant antibodies at 4ºC for 12-h followed by incubation with protein A+G sepharose beads for 5-h. After 3x washes with wash buffer, beads were boiled at 95˚C for 10-min. After boiling, supernatant was used for immunoblotting along with the input. After electrophoresis

SDS-PAGE, lysate proteins were transferred to a nitrocellulose membrane and immunoblotting was performed as described previously (Ekka et al, 2020; Kumar et al, 2017) using specific antibodies and blots were developed using SuperSignal® West Pico or Femto chemiluminescence Substrate (Thermo Scientific) by following manufacturer's instructions.

**Generation of pS33-HP1 antisera**

Custom antibodies were raised against HP1 phosphorylated at S33 (anti-pS33-PfHP1) or an unphosphorylated version of the S33 (HP1 non-phospho) by M/s Antagene Inc. For phospho antibody generation, a peptide with a sequence: Cys-YLVKWKGY(p)SDDENTW and for non-phospho peptide Cys-YLVKWKGYSDDENTW was used for immunization. Phospho-antibodies were purified by affinity chromatography using these peptides. For Western blotting, 1:100-200 dilution of the purified phospho and non-phospho antibody was used and for IFAs 1:25 of anti-pS33HP1 was used. Total HP1 antisera were raised in rabbit against bacterially expressed HP1 (a gift from Dr. Krishanpal Karmodiya, IISER Pune).

**SUrfaceSEnsing of Translation (SUnSET) assay or Puromycin incorporation assay**

Protein synthesis was assessed using a SUnSET assay (Hobson et al, 2020; McLean & Jacobs-Lorena, 2017) for which parasites were treated with 1µM puromycin (structural analog of tyrosyl-tRNA) for 1-h. Parasites were harvested and protein lysates were prepared using 2% SDS, which were subjected to Western blot analysis. To ensure equal loading of proteins, the PVDF membrane was stained with Ponceau S before blocking with 3% BSA for 1-h. After blocking, blots were incubated with anti-puromycin antibody (Cat no. MABE343,

Merck) overnight at 4ºC. Subsequently, blots were washed with 1x TBST and incubated with α-FCY-subclass 2a mouse IgG antibody for 1-h. The blots were developed by Pico or Femto chemiluminescence substrate from Pierce (USA) after exposure of the membrane to X-ray films.

### Quantitative RT-PCR

RNA isolation was carried out using TRIzol reagent (G Biosciences). Real time PCR was carried out using CFX96 Real Time PCR detection system (Biorad). Quantification of the expression was done by determining SYBR green incorporation in the amplicons (G Biosciences). For qRT-PCR, 1 μg of DNAse free RNA was used for cDNA synthesis using Verso cDNA synthesis kit (Thermo scientific), as per the manufacturer's recommendation. Random hexamers supplied with the kit were used for the cDNA synthesis. 18srRNA and Actin were used as a housekeeping gene for normalization. A typical PCR reaction conditions were: 95°C for 3-min, 95°C for 15-sec, 60°C for 1-min and 40 cycles. Finally, the relative fold change in gene expression was determined as $2^{-\Delta\Delta Ct}$. Primers details for the qRT-PCR are given in (Supplementary Table S2).

### RNA sequencing

Nf54 (control) and PfPPM2-HA-glmS$^{Nf54}$ lines were treated with GlcN and were harvested at first cycle at the schizont stage (42-44 h.p.i) in triplicate. RNA isolation was performed using the phenol-chloroform extraction method, followed by quantification using a Qubit fluorometer (Thermo Fisher Scientific). For all samples, 1μg of RNA was used as an input for mRNA enrichment using the NEBNext Poly (A) mRNA magnetic isolation module (New

England Biolabs #E7490S) according to the manufacturer's instructions. A NEBNext Ultra II RNA library prep kit for Illumina (New England Biolabs #E7770S) was used to prepare the library according to the manufacturer's instructions, and quality checks were performed using Agilent TapeStation. Paired-end sequencing was performed on an Illumina Nextseq550 platform (75 bp read length).

**RNAseq Data Analysis**

RNA-seq data analysis was performed using *in-house* scripts and pipeline. The quality of sequenced samples was accessed using FASTQC (http://www.bioinformatics.babraham.ac.uk/projects/fastqc) and adapters were trimmed with Trim Galore ( https://doi.org/10.5281/zenodo.5127899) and cutadapt (https://doi.org/10.14806/ej.17.1.200). High quality trimmed reads were aligned in paired-end mode to *Plasmodium falciparum* 3D7 genome ver. 66 using HISAT2 (Kim et al, 2019) with default settings. Differential gene expression analysis was performed using DEseq2 (Love et al, 2014) using standard analysis protocol. Gene set enrichment analysis was performed using ClusterProfiler (Yu et al, 2012) and results were plotted with custom R and python scripts.

**Chromatin Immunoprecipitation (ChIP) and ChIP-qPCR**

Parasites cultured in the presence (2.5 mM GlcN) or absence of glucosamine (GlcN) and were harvested at the end of first cycle at ~90 h.p.i by fixing the cells. The iRBCs were cross linked using 1% formaldehyde (Thermo Scientific, 28908) for 10-min. Subsequently, 150 mM glycine was then added for quenching the cross-linking reaction for 10-min at ambient temperature followed by centrifugation at 6000 rpm at 4˚C for 5-min. Pellet was washed three times using 1xcold PBS (supplemented with protease inhibitors) before proceeding for lysis. Sample homogenization was performed using swelling buffer (25 mM Tris pH 7.9, 1.5 mM MgCl2, 10 mM KCl, 0.1% NP 40, 1 mM DTT, 0.5 mM PMSF, 1 × PIC) followed by cell lysis in sonication buffer (10 mM Tris–HCl pH 7.5, 200 mM NaCl, 1% SDS, 4% NP-40,1 mM PMSF, 1X protease inhibitor cocktail). First, pre-clearing was performed for 2-h at 4°C by incubating the chromatin fraction with- protein A conjugated sepharose beads (G-Biosciences) with continuous gentle mixing. Pre-cleared lysate was incubated with the relevant antibody for 12-h at 4°C followed by incubation with saturated Protein A Sepharose beads for 4-h at 4 °C. Bound chromatin was finally washed with low salt wash buffer, high salt wash buffer, LiCl wash buffer, TE wash buffer and eluted using ChIP elution buffer (1% SDS, 0.1 M sodium bicarbonate). Both IP sample and input were reverse cross-linked using 0.3 M NaCl overnight at 65°C along with RNAase followed by Proteinase K treatment at 42°C for 1-h. Finally, bound DNA was purified using phenol-chloroform precipitation. The occupancy of genes of interest was assessed by ChIP-qPCR using the SYBR Green Master Mix (G Biosciences). The amount of template used for qPCR was 1 µl of 1% input DNA and 2 µl of undiluted ChIP DNA. qPCR was performed using previously described primers corresponding to AP2-G and Pfs16 locus (Table S2) (Brancucci et al, 2014). The amount of DNA amplified was compared to the input and % recovery of bound DNA was determined.

**Phosphoproteomics and Mass spectrometry**

***Proteomics Studies***

*Cell lysis and protein extraction and TMT labeling*

The 3D7 or PfPPM2-HA-glmS$^{3D7}$ parasites were cultured and synchronized using sorbitol. Ring stage parasites were treated with glucosamine for 72-h. The parasites were subsequently harvested upon maturation to schizonts in cycle 1 (~44 hours post-infection), by lysing the infected red blood cells with saponin. The parasite pellet was washed and dissolved in protein extraction buffer (50 mM triethylammonium bicarbonate (TEAB), 2% SDS, and 1X protease and phosphatase inhibitors) followed by sonication and centrifugation. Protein estimation was done by Bicinchoninic Acid (BCA) assay. Equal amounts of protein were taken from all sets of parasites and subjected to reduction with 20 mM Dithiothreitol (DTT) at 60°C for 30-min, followed by alkylation with 20 mM Iodoacetamide (IAA) for 10-min in the dark at ambient temperature. Subsequently, proteins were digested using trypsin and Lys-C mix (Promega, Madison, USA) used at a ratio of 100:1. Tryptic peptides were cleaned using a C18 column and dried with a SpeedVac concentrator (Thermo Fisher Scientific, USA). The peptides were quantified using the Pierce Quantitative Colorimetric Peptide Assay Kit.Further, 500 µg of peptides from each sample were processed for TMT labelling following manufacturer's instructions (Thermo Scientific). Labelling efficiency of >95% was achieved and the reaction was stopped by adding 8 µl of hydroxylamine. The labelled peptides were pooled, dried, and cleaned using C18 Sep-Pak cartridges (Waters, Milford, MA, USA).

*Phosphopeptide enrichment and fractionation*

50µg of the peptide sample was used for total proteome analysis, and the remaining sample was subjected to phosphopeptide enrichment. For the initial phosphopeptide enrichment, the TiO2 Phosphopeptide Enrichment Kit was utilized (Thermo Scientific). The subsequent enrichment step was performed on the collected flow-through using the High-Select Fe-NTA Phosphopeptide Enrichment Kit (Thermo Scientific). The eluents from both enrichment steps were combined and subjected to fractionation using C18 Stage Tips. A total of 24 fractions were collected and then concatenated into six fractions. The peptide samples were dried and dissolved in 0.1% formic acid prior to mass spectrometry analysis.

*LC-MS/MS analysis by Data-Dependent Acquisition*

Mass spectrometry analysis was performed on an Orbitrap Fusion Tribrid Mass Spectrometer linked to an Easy-nLC 1200 nano-flow UPLC system (Thermo Fisher Scientific, Germany). The peptide fractions were loaded on nanoViper column (2 cm, 3 µm C18 Aq) (Thermo Fisher Scientific) and were then resolved on an analytical column (75 µm × 15 cm, C18, 2 µm particle size) (Thermo Fisher Scientific). The peptides were separated by passing solvent A (0.1% formic acid) and solvent B (80% acetonitrile in 0.1% formic acid) using a gradient mode over a 120-min at a flow rate of 300 nl/min.

Data-dependent acquisition (DDA) mode was used for data acquisition. Precursor ions were acquired in full MS scan mode in the range of 400-1600 m/z, using an Orbitrap mass analyzer resolution of 120,000 at 200 m/z. An automatic gain control (AGC) target value of 2e5 with an injection time of 55ms and dynamic exclusion of 30 seconds was used. The selection of the most intense precursor ions was performed at top speed DDA mode, selected using a quadrupole with an isolation window of 2 m/z. The filtered precursor ions were subsequently fragmented using higher-energy collision-induced dissociation (HCD) with 32±3% normalized collision energy and MS/MS scans in the range of 110-2000 m/z were acquired by using Orbitrap mass analyzer with 30,000 mass resolution at 200 m/z. AGC target was set to 1e5 with an injection time of 200ms. For each fraction, MS/MS data was acquired in duplicates.

*Data analysis*

MS/MS raw data was used to search against a combined protein database of *P. falciparum* 3D7 (downloaded from PlasmoDB web resource, version 46) and *Homo sapiens (RefSeq v92)* using SEQUEST and Mascot (version 2.4.1) via Proteome Discoverer v2.2 (Thermo Fisher Scientific) as described previously (Rawat et al, 2023). A fold change cut-off of 1.5 with a p-value ≤ 0.05 was used to identify significantly altered phosphorylated proteins. These proteins were then analyzed for the enrichment of biological processes using Gene Ontology (GO) analysis through PlasmoDB.

**Densitometry and statistical analysis**

Image J (NIH) software was used to perform densitometry of Western blots. The band intensity of the loading control was used for normalization. Statistical analysis was performed using Prism (Graph Pad software Inc USA). Data is represented as mean ± Standard error of mean (SEM), unless indicated otherwise and $p < 0.05$ was considered as statistically significant.

**Table-S1: A selected list of key putative targets of PfPM2 that may be related to its role in parasite division and sexual conversion.**

Significantly altered Hyper or Hypo-phosphorylated proteins upon PfPPM2 depletion (Fig. 2A, Dataset S1) belonging to families that may be related to PfPPM2 function

| Phosphorylated proteins | PlasmoDB ID | Phosphosites |
|---|---|---|
| **Chromatin remodelling proteins and epigenetic regulators** | | |
| ISWI chromatin-remodeling complex ATPase<br>Histone H3 variant (H3.3)<br>Histone H3<br>AP2 domain transcription factor, putative (AP2- P)<br>Heterochromatin protein 1 | PF3D7_0624600<br>PF3D7_0617900<br>PF3D7_0610400<br>PF3D7_1107800<br>PF3D7_1220900 | S1895,S1898<br>S28<br>S32<br>S1241; S1244<br>S33 |
| **Translational machinery** | | |
| Eukaryotic translation initiation factor 2-alpha kinase<br>Elongation factor 1-alpha | PF3D7_0628200<br>PF3D7_1357000 | S1973,S1773<br>T213 |
| **IMC complex** | | |
| Glideosome-associated protein 40, putative | PF3D7_0515700 | S420 |
| **Chromatin remodelling and assembly** | | |
| 14-3-3 protein<br>Bromodomain protein 2, putative<br>MORC family protein | PF3D7_0818200<br>PF3D7_1212900<br>PF3D7_1468100 | S256<br>S408<br>S2401 |
| **Basal complex** | | |
| Protein CINCH<br>Schizont egress antigen-1<br>Basal complex transmembrane protein 2<br>Basal complex transmembrane protein 1 | PF3D7_0407800<br>PF3D7_1021800<br>PF3D7_0704100<br>PF3D7_0611600 | S1561<br>T1232,S1234,S1237<br>S782, S787<br>S434 |

**Table-S2: List of all primers used in this study:**

| Purpose | Forward (5'-3') | Reverse (5'-3') |
|---|---|---|
| *PfPPM2-HA-GlmS*: 3' Homology arm cloning | 1 AGAGATCTAATGATGAAGAAACAGGAGAAGATG | 2 AGCTGCAGCTTCAATGTCGTCGATATTAAGAAATTTC |
| *PfPPM2-HA-GlmS*: 5' integration genotyping | 3 AAAATAGGAATAATGGAGATGAAG | 4 ATTTTCTTCTACATCTCCACATG |
| *PfPPM2-HA-GlmS* : 3'integration genotyping | 5 ATAACAATTTCACACAGGAAACAG | 6 CATCTGTATTATATGCATAAATGTTTC |
| *PfPPM2-HA-GlmS*: WT locus Genotyping | 3 AAAATAGGAATAATGGAGATGAAG | 6 CATCTGTATTATATGCATAAATGTTTC |
| For amplificationof 2A Skip-yDHOH locusfrom pSLI-N-sandwich loxP vector | 7 CGGGGTACCGACTATAAGGACCACGACGGAGACTACAAGGATCATGATATTGATTACAAAGACGATGACGATAAGGCTAGCGGAGAAGGAAGAGGAAGT | 8 CCGCTCGAGAATGCTGTTCAACTTCCCAC |
| Cloning of HP1(WT)HR in 3x Flag-yDHODH-BSD vector | 9 GGACTAGTATGGAAAGGATATTCAGATGATGAG | 10 CGGGGTACCAGCTGTACGGTATCTTAGTCTTG |
| Cloning of HP1(S33A)HR in 3x Flag-yDHODH-BSD vector | 11 GGACTAGTATGGAAAGGATATGCTGATGATGAG | 10 CGGGGTACCAGCTGTACGGTATCTTAGTCTTG |
| *PfPPM2-3xHA-GlmS*$^{NF54:}$ $_{HP1/HP1\text{-}S33A\text{-}Flag}$: 5' integration genotyping | 12 ATGAAGAATTTGAAATTGGTG | 13 GTTTAATACATAGGACAAATAGT |
| *PfPPM2-3xHA-GlmS*$^{NF54:}$ $_{HP1/HP1\text{-}S33A\text{-}Flag}$: WT locus genotyping | 12 ATGAAGAATTTGAAATTGGTG | 15 TTCCAATTGATATTATTATTACC |
| For amplificationof HP1 gene | 16 CGGGGTACCATGACAGG | 17 TTCTCCTTTCTGCAGAG |

| | | |
|---|---|---|
| from 3D7 gDNA | TTCAGATGAAGAATTTG | CTGTACGGTATCTTAGT CTTG<br>17'<br>GTGCCTAGGAGCTGTA CGGTATCTTAGTCTTG |
| For apmplification of GFP from cpARL-BSD vector | 18<br>CTGCAGAAAGGAGAAGA AC | 19<br>TCCCATTTTGAGCTCTT TGTATAGTTCATCCATG CCATG |
| For amplificationof DD domain from pSLI-DD vector | 20<br>GAGCTCAAAATGGGAGT GCAG | 21<br>GTGCCTAGGTTAAGGTT CCGGTTTTAGAAGCTC |
| Overlapping PCR primers for HP1-GFP | 16<br>CGGGGTACCATGACAGG TTCAGATGAAGAATTTG | 19<br>TCCCATTTTGAGCTCTT TGTATAGTTCATCCATG CCATG |
| Overlapping PCR primers for GFP-DD | 18<br>CTGCAGAAAGGAGAAGA AC | 21<br>GTGCCTAGGTTAAGGTT CCGGTTTTAGAAGCTC |
| Overlapping PCR primers for HP1-GFP-DD domain | 16<br>CGGGGTACCATGACAGG TTCAGATGAAGAATTTG | 21<br>GTGCCTAGGTTAAGGTT CCGGTTTTAGAAGCTC |
| SDM primers for HP1 S33A for cloning in cpARL-GFP-DD vector | 22<br>AAATGGAAAGGATATGC TGATGATGAGAATACT | 23<br>AGTATTCTCATCATCAG CATATCCTTTCCATTT |
| SDM primers for HP1 S33D for cloning in cpARL-GFP-DD vector | 24<br>AAATGGAAAGGATATGA TGATGATGAGAATACT | 25<br>AGTATTCTCATCATCAT CATATCCTTTCCATTT |
| **Real Time PCR Primers used in the study** | | |
| 18SRNA | 26<br>GCTGACTACGTCCCTGC CC | 27<br>ACAATTCATCATATCTT TCAATCGGTA |
| actin | 28<br>AGCAGCAGGAATCCACA CA | 29<br>TGATGGTGCAAGGGTT GTAA |
| hp1 | 30<br>CGAAAGCTAATGAGACA AATGGT | 31<br>CGTCGGGGTGCTAAGG AAC |
| ap2-g | 32<br>TGGTGGTAATAAGAACA ACAGAGGT | 33<br>CCATCATAATCTTCTTC TTCGTCG |

| | | |
|---|---|---|
| *Pfs16* | 34 TTCTTCGCTTTTGCAAACCT | 35 AGCATGAAGAGAGGCACCTG |
| **ChIP-qPCR Primers used in the study** | | |
| pfap2-g | 36 TGGTGGTAATAAGAACAACAGAGGT | 37 CCATCATAATCTTCTTCTTCGTCG |
| Pfs16 | 38 AGTTCTTCAGGTGCCTCTCTTCA | 39 AGCTAGCTGAGTTTCTAAAGGCA |
| Actin | 40 AGCAGCAGGAATCCACACA | 41 TGATGGTGCAAGGGTTGTAA |

**Table-S3: List of antibodies used in the study**

| Antigen | Source | | Dilution used | |
|---|---|---|---|---|
| | | | WB | IFA |
| Centrin-1 | Mouse | (20H5 monoclonal mice antibody);Sigma | NA | 1:200 |
| Alpha-tubulin | Mouse | monoclonal, clone DM1A; Sigma | NA | 1:200 |
| HA-Tag | Mouse | Sigma | 1:1000 | 1:200 |
| Puromycin | Mouse | Sigma | 1:1000 | NA |
| Flag-Tag | Mouse | Cat. No.- sc-51590 | 1:1000 | 1:200 |
| GFP | Mouse | Sigma | 1:1000 | 1:200 |
| β-Actin | HRP conjugated | Santacruz | 1:1000 | NA |
| BiP (MRA-1246) | Rabbit | Bei resources | 1:1000 | NA |
| HA-Tag (C29F4 mAb) | Rabbit | Cell signaling and technology | 1:1000 | 1:200 |
| Tri-Methyl-Histone H3 (Lys9) | Rabbit | Cell signaling and technology | 1:1000 | 1:200 |
| Histone H3 (D2B12) | Rabbit | Cell signaling and technology | 1:1000 | 1:200 |
| H3S10ph (ab5176) | Rabbit | Abcam | 1:1000 | NA |
| H3S28ph (ab5169) | Rabbit | Abcam | 1:1000 | NA |
| Phospho-EIF2S1 (Ser51) | Rabbit | Thermo | 1:1000 | NA |
| eIF2A- D7D3 (Total eIF2α) | Rabbit | Thermo | 1:1000 | NA |
| PfIMC1g | Mouse | Generated in Dr. Sharma laboratory | NA | 1:200 |
| Anti-PfMSP1 | Rabbit | Bei resources | NA | 1:200 |
| Pfs16 (MRA1276) | Rabbit | Bei resources | NA | 1:200 |
| Pfs230 (MRA-878A) | Rabbit | Bei resources | NA | 1:200 |
| pS33-HP1 and non-phospho HP1 | Rabbit | Antagene | 1:100 | 1:25 |
| HP1 (total) | Rabbit | Generated in Dr. Sharma laboratory | NA | 1:200 |

**Supplementary figures**

**Supp. Fig. S1: Generation of** PfPPM2-HA-glmS$^{3D7}$/NF54 **parasites for conditional knockdown of PfPPM2.**

A. Schematic representation of the strategy used to modify the *PfPPM2* locus by single-crossover homologous recombination and selection-linked integration (SLI). A homology region corresponding to the 3'-end of *PfPPM2* was cloned in pGlms-SLI vector and resultant construct was used for transfection.

B and C. PfPPM2-HA-GlmS-SLI construct was transfected in either 3D7 (B) or NF54 (C) parasites and subjected to drug selection. Parasite clones were obtained in both cases which were confirmed by genotyping PCR performed using primers indicated in Panel A that yielded amplicons of the expected size indicated in the table.

D. Western blotting of PfPPM2-HA-glmS$^{NF54}$ parasites, which were either left untreated or treated with GlcN (cycle 0, rings) and parasite lysates were prepared (schizonts, cycle 1) using anti-HA antibody. Anti-BiP antibody was also used to probe the blots.

**Supp. Fig. S2: Effect of GlcN tretament on NF54 and 3D7 parasites**

The unmodified lines 3D7 and NF54 parasites were synchronized and ring stage parasites and were either left untreated or treated with 2.5mM GlcN for one cycle as described for with PfPPM2-HA-glmS$^{NF54/3D7}$ (Fig. 1C). Parasites were analyzed at indicated time points by flow cytometry and % parasitemia was determined. Data shown in the line graph are a representative of three independent biological replicates. *Inset*, fold change in growth of 3D7 and NF54 parasites cultured in the presence or absence of GlcN from three independent biological replicates (SEM ± SE, n=3, ANOVA, P > 0.05, ns-non-significant).

**Supp. Fig. S3 Effect of PfPPM2 depletion on parasite development**

A. 3D7 and PfPPM2-HA-glmS$^{3D7}$ parasites were synchronized and ring stage parasites were either left untreated or treated with 2.5mM GlcN. The individual parasitic stages-rings, trophozoites, and schizonts were counted at the indicated time points from Giemsa-stained thin blood smears of parasite cultures and are represented as % of total number of parasites. There was no significant difference in intraerythrocytic development in GlcN treated and untreated parasites. Data representative of three independent replicates is provided (Mean ± SEM, ANOVA, N=3, P > 0.05, ns-non-significant).

B. Representative images of Giemsa-stained thin blood smears of parasite culture (panel A) when parasites were either left untreated or treated with 2.5mM GlcN. There was no significant difference in intraerythrocytic development of parasite upon PfPPM2 depletion as quantitated in the above panel.

C. PfPPM2-HA-glmS$^{3D7}$ schizonts were cultured in the presence or absence of glucosamine (GlcN) and incubated with fresh erythrocytes. % schizont or ring infected erythrocytes was determined after 10-h. PfPPM2-depleted parasites do not show any significant difference in the number of new rings formed (SEM ± SE, n=3, ANOVA; P>0.05, ns-non-significant).

D. PfPPM2-HA-glmS$^{3D7}$ parasites were treated with GlcN as described in Fig. 1B and C. Thin blood smears were made ~40-44 hpi in cycle 1. IFA was performed using IMC1g and MSP1 antibodies. Scale bar =1 μm. (SEM ± SE, n=3,t-test; *P<0.05).

**Supp. Fig. S4 Work flow for phosphoproteomics studies**

PfPPM2-HA-glms$^{3D7}$ parasites were synchronized to obtain rings, which were treated with GlcN and parasites were harvested at schizont stage in cycle 1 to perform comparative phosphoproteomic and proteomic analyses as indicated in the schematic (See Methods for details).

**Supp. Fig. S5: PfPPM2 regulates protein synthesis**

A. eIF2α kinase PfPK4 was hyper phosphorylated at two sites (*) that are upstream of its kinase domain (KD).

B. Schematic represents the role of PK4 in regulating eIF2α. An upstream signal activates PfPK4, which subsequently phosphorylates eIF2α at S51 (S59 in P. falciparum), leading to its deactivation resulting in translational repression and inhibition of protein synthesis (Zhang et al, 2017).

C. Western blotting of lysates from PfPPM2-HA-glmS$^{3D7}$ parasites, which were either left untreated or treated with GlcN using antibodies against phospho-eIF2α or total eIF2α. Anti-BiP antibody was used to probe the blots as a loading control. *Lower panel,* fold change in eIF2α phosphorylation upon PfPPM2 depletion in experiments described in the upper panel was determined by densitometry of eIF2α bands in the Western blots, which was normalized with total eIF2α (SEM ± SE, n=3, ANOVA, *P<0.05,P > 0.05,ns).

D. PfPPM2-HA-glmS$^{3D7}$ parasites were synchronized and were grown in the presence or absence of GlcN. At schizont stage (42hpi, cycle 1), puromycin was added for 1-h and parasites were harvested for Western blotting, which was performed using an antibody against puromycin to assess protein synthesis in the parasite. *Right Panel,* Ponceau S-stained membrane which was used for Western blotting.

**Supp. Fig. S6 HP1 overexpression reverts defects in parasite development**

A. Western blotting of PPM2-HA-GlmS$^{NF54}$ parasites or HP1-GFP overexpressing (*PfPPM2-HA-glmS$^{Nf54:}$HP1-GFP*$^{OE}$ parasites, which were either left untreated or treated with GlcN (cycle 0, rings) and parasite lysates were prepared (schizonts, cycle 1) using anti-GFP antibody. β-actin antibody was also used to probe the blots.

B. *PfPPM2-HA-glmS$^{Nf54:}$HP1-GFP*$^{OE}$ parasites were used for IFA performed with anti-HA and anti-GFP antibodies which revealed partial co-localization between HP1 and PfPPM2.

C. Gametocyte formation was induced in GlcN treated or untreated PPM2-HA-glmS$^{NF54}$ and *PfPPM2-HA-glmS$^{Nf54:}$HP1-GFP*$^{OE}$ parasites lines. % gametocytaemia was determined 7 days post induction.

**Supp. Fig. S7. Generation of** PfPPM2-HA-GlmS$^{Nf54:HP1/S33A-Flag}$ parasites

A. Schematic representation of the strategy used to modify the *HP1* locus to introduce a Flag tag at its C-terminus in PfPPM2-HA-glms$^{NF54}$ background using selection-linked integration (SLI). Another parasite line in which S33A mutation was made in HP1 along with a Flag tag was also generated (HP1-S33A-flag). A homology region corresponding to the 3'-end to WT HP1 or a version with S33A mutation was cloned to generate the targeting construct PfPPM2-HA-GlmS$^{Nf54:HP1/S33A-Flag}$.

B. Plasmid construct described in panel A was transfected in PfPPM2-HA-glms$^{NF54}$ parasites and subjected to drug selection. Parasites (PfPPM2-HA-GlmS$^{Nf54:HP1/S33A-Flag}$) obtained after drug selection were subjected to genotyping using PCR primers indicated in Panel A that yielded amplicons of the expected size indicated in the table and exhibited the presence of both unmodified (WT) HP1 as well as modified (HP1/S33A-Flag) parasites.

*Right panel*, sequencing of PCR product spanning the region coding S33 in PfPPM2-HA-GlmS:HP1/S33A-Flag parasites confirmed the mutation of S33 to A in PfPPM2-HA-GlmS$^{Nf54:HP1\text{-}S33A\text{-}Flag}$ parasites.

**Supp. Fig. S8 Conditional overexpression of HP1/S33A/S33D**

A. Schematic representing HP1 or its S33A/D mutants tagged with GFP and DD domain at C-terminus which was used for overexpression in NF54 parasites.

B. HP1/S33A/S33D-GFP-DD parasites were synchronized and ring stage parasites were cultured in the presence or absence of Shld-1 for 72-h. Western blotting using anti-GFP antibodies revealed successful expression of these proteins (HP1/S33A/S33D-GFP-DD) in the presence of Shld-1 but a small amount of proteins was also expressed in its absence and was indicative of "leaky" expression.

C. IFA was performed on thin blood smears of HP1/S33A/S33D-GFP-DD parasites cultured in the presence of Shld1 using anti-H3K9me3 or GFP antibodies, which revealed reduced H3K9me3 staining in S33A parasites.

**Supp. Fig. S9: RNAseq analysis to identify changes expression of genes upon PfPPM2 depletion**

A. PfPPM2-HA-glms$^{Nf54}$ parasites were treated with GlcN as described in Fig. 1B and C. RNAseq analysis was performed to compare global changes in transcriptome of these parasites upon PfPPM2 depletion. The volcano plot illustrates the differentially expressed genes, with some of the significantly altered key genes are labelled (Supplementary Data Set S3).

B. Gene Ontology terms describing biological processes significantly altered in GSEA upon PfPPM2 depletion (Supplementary Data Set S3).

**Supp. Fig. S10 pS33-HP1 during sexual conversion**

Sexual conversion of an unrelated *P. falciparum* strain was induced by serum depletion on an unrelated parasite line. IFA was performed on parasites pre- and 3 days post-induction (Fig. 6B) using anti-pS33-HP1 and anti-Pfs230 antibodies. HP1-S33-P signal was present mainly in uninduced schizonts whereas it was reduced post induction and it was almost completely absent from gametocytes that were stained with anti-Pfs230. (SEM ± SE, n=2, -t-test, *P<0.05).

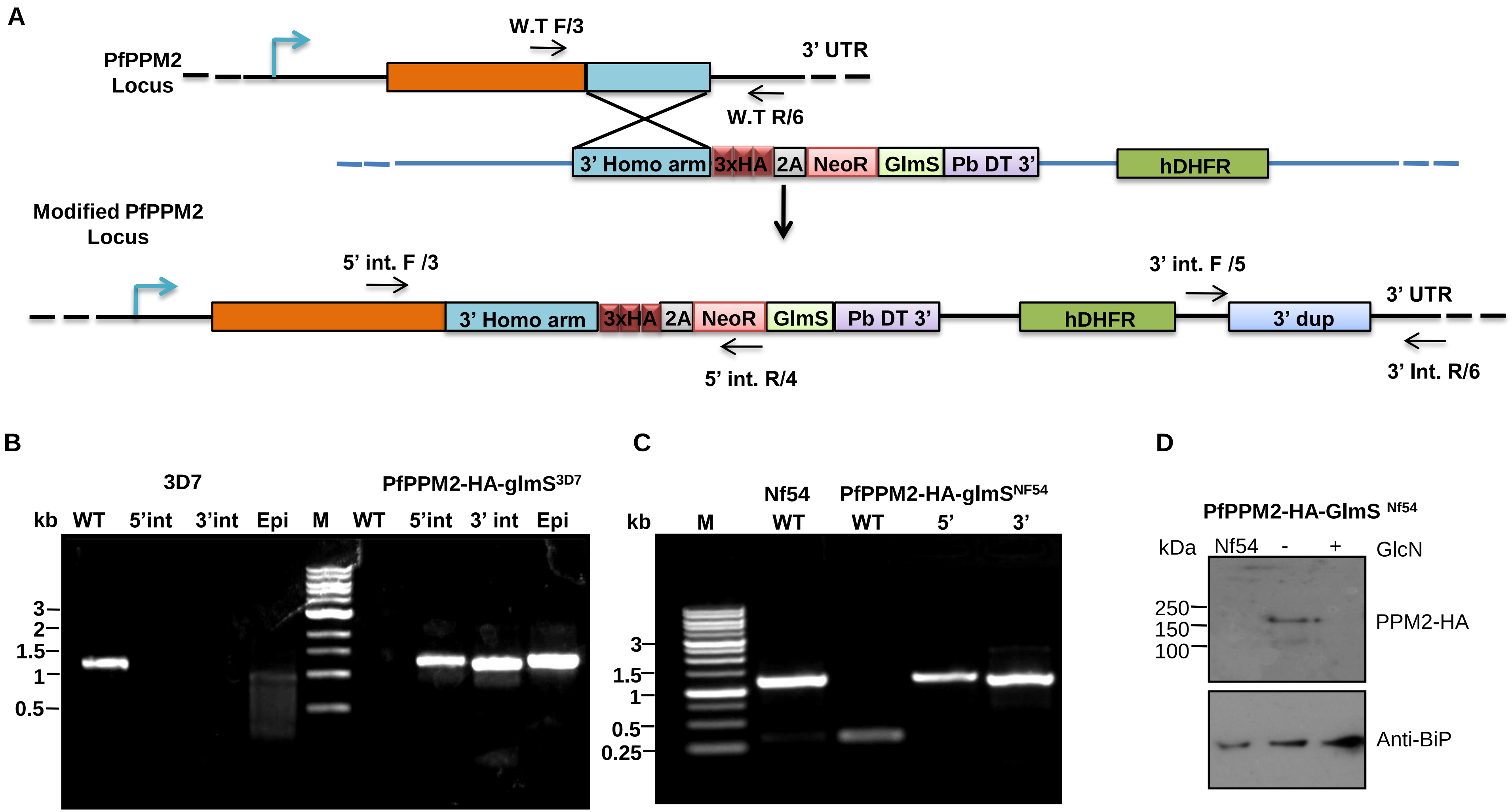


| Locus | Primer set | Expected size |
|---|---|---|
| WT | 3/6 | 1.2 kb |
| 5' integration | 3/4 | 1.2 kb |
| 3' integration | 5/6 | 1.1 kb |

**Fig. S1**

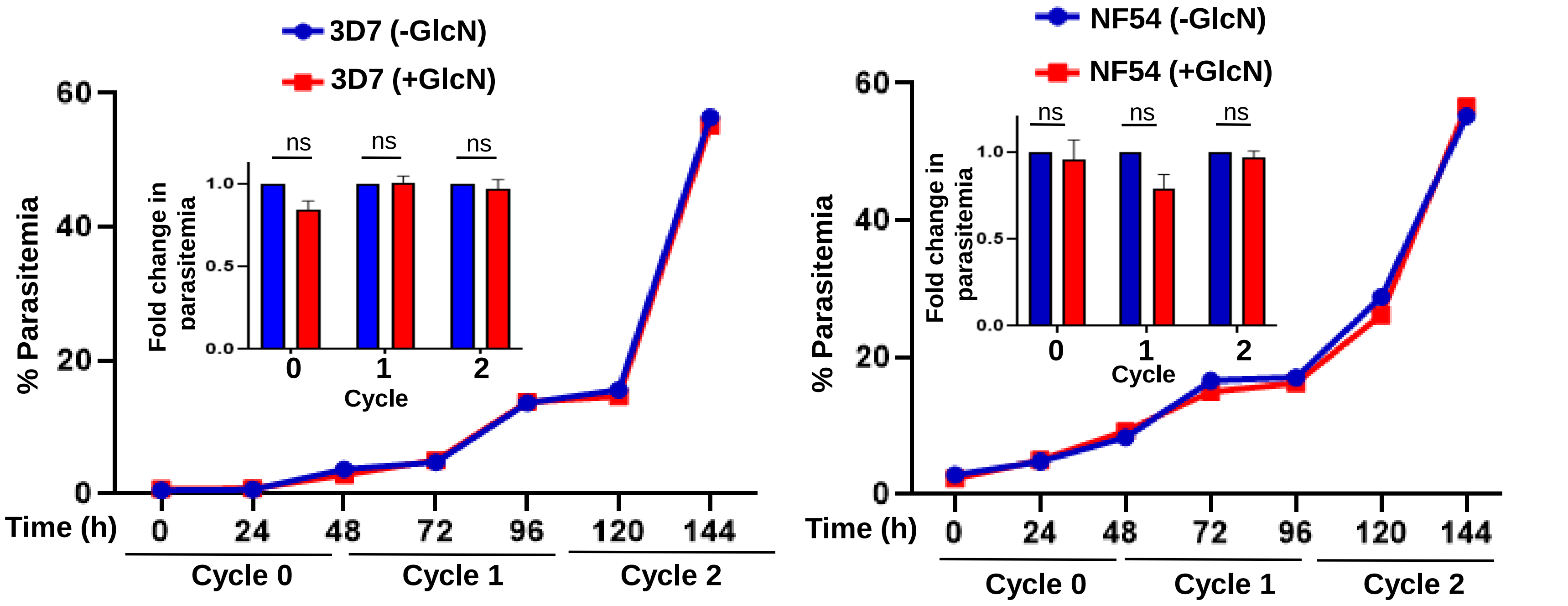
3D7 (-GlcN)
3D7 (+GlcN)
% Parasitemia
60
40
20
0
Time (h)
0
24
48
72
96
120
144
Cycle 0
Cycle 1
Cycle 2
Fold change in parasitemia
1.0
0.5
0.0
ns
ns
ns
0
1
2
Cycle
NF54 (-GlcN)
NF54 (+GlcN)
% Parasitemia
60
40
20
0
Time (h)
0
24
48
72
96
120
144
Cycle 0
Cycle 1
Cycle 2
Fold change in parasitemia
1.0
0.5
0.0
ns
ns
ns
0
1
2
Cycle

Fig. S2

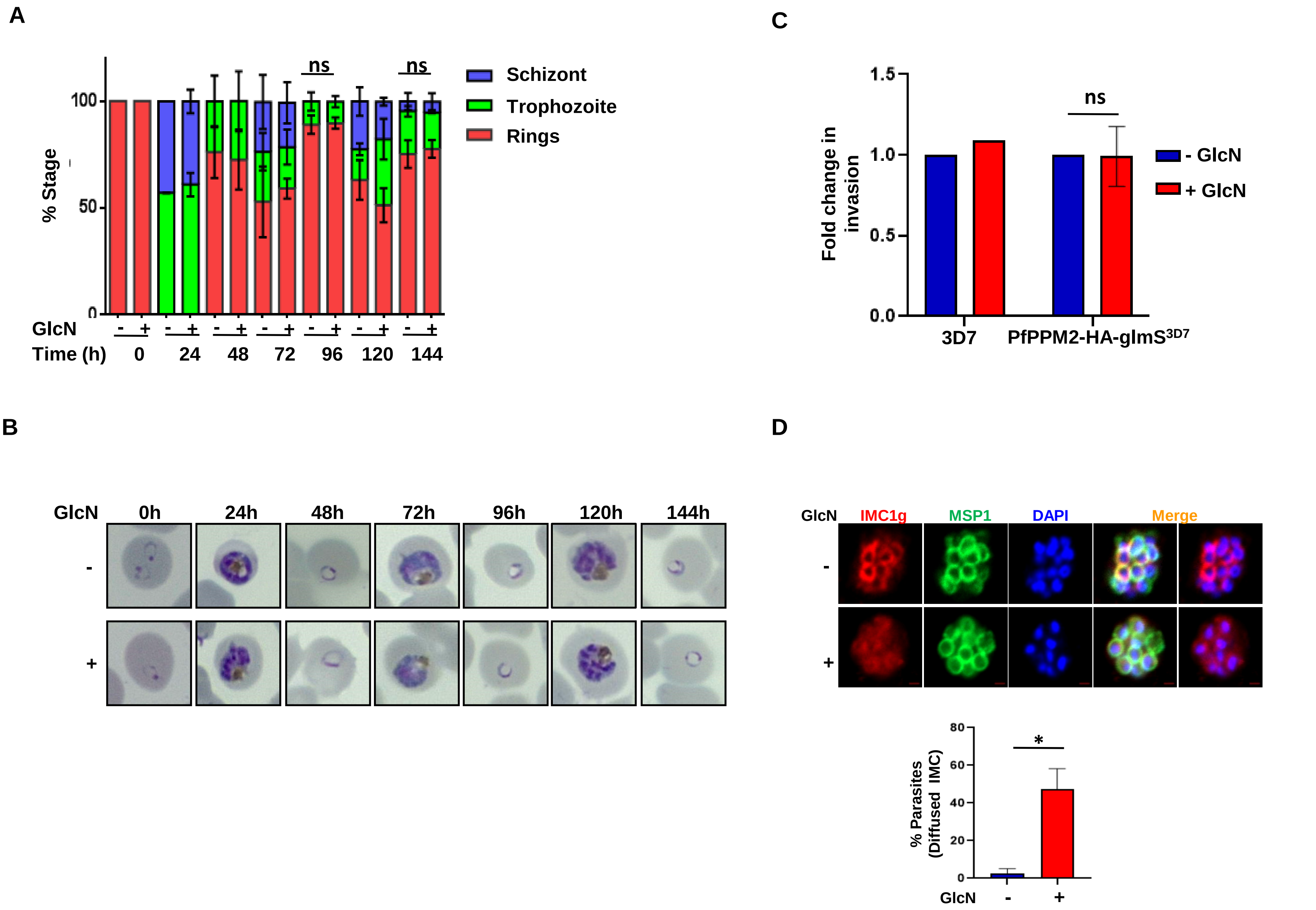
A
% Stage
100
50
0
ns
ns
Schizont
Trophozoite
Rings
GlcN
- + - + - + - + - + - + - +
Time (h) 0 24 48 72 96 120 144
B
GlcN 0h 24h 48h 72h 96h 120h 144h
-
+
C
Fold change in invasion
1.5
1.0
0.5
0.0
ns
- GlcN
+ GlcN
3D7
PfPPM2-HA-glmS3D7
D
GlcN IMC1g MSP1 DAPI Merge
-
+
% Parasites (Diffused IMC)
80
60
40
20
0
*
GlcN - +

Fig. S3

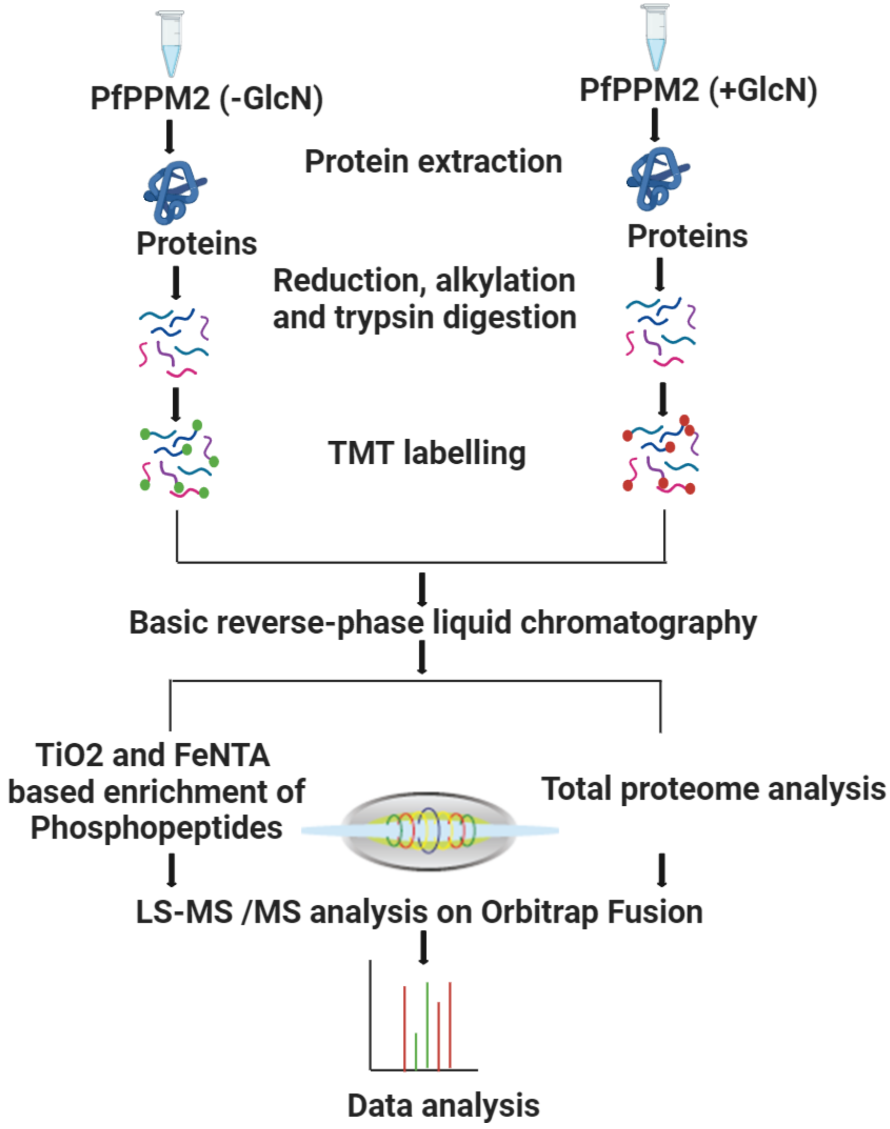
PfPPM2 (-GlcN)
PfPPM2 (+GlcN)
Protein extraction
Proteins
Proteins
Reduction, alkylation
and trypsin digestion
TMT labelling
Basic reverse-phase liquid chromatography
TiO2 and FeNTA
based enrichment of
Phosphopeptides
Total proteome analysis
LS-MS /MS analysis on Orbitrap Fusion
Data analysis

**Fig. S4**

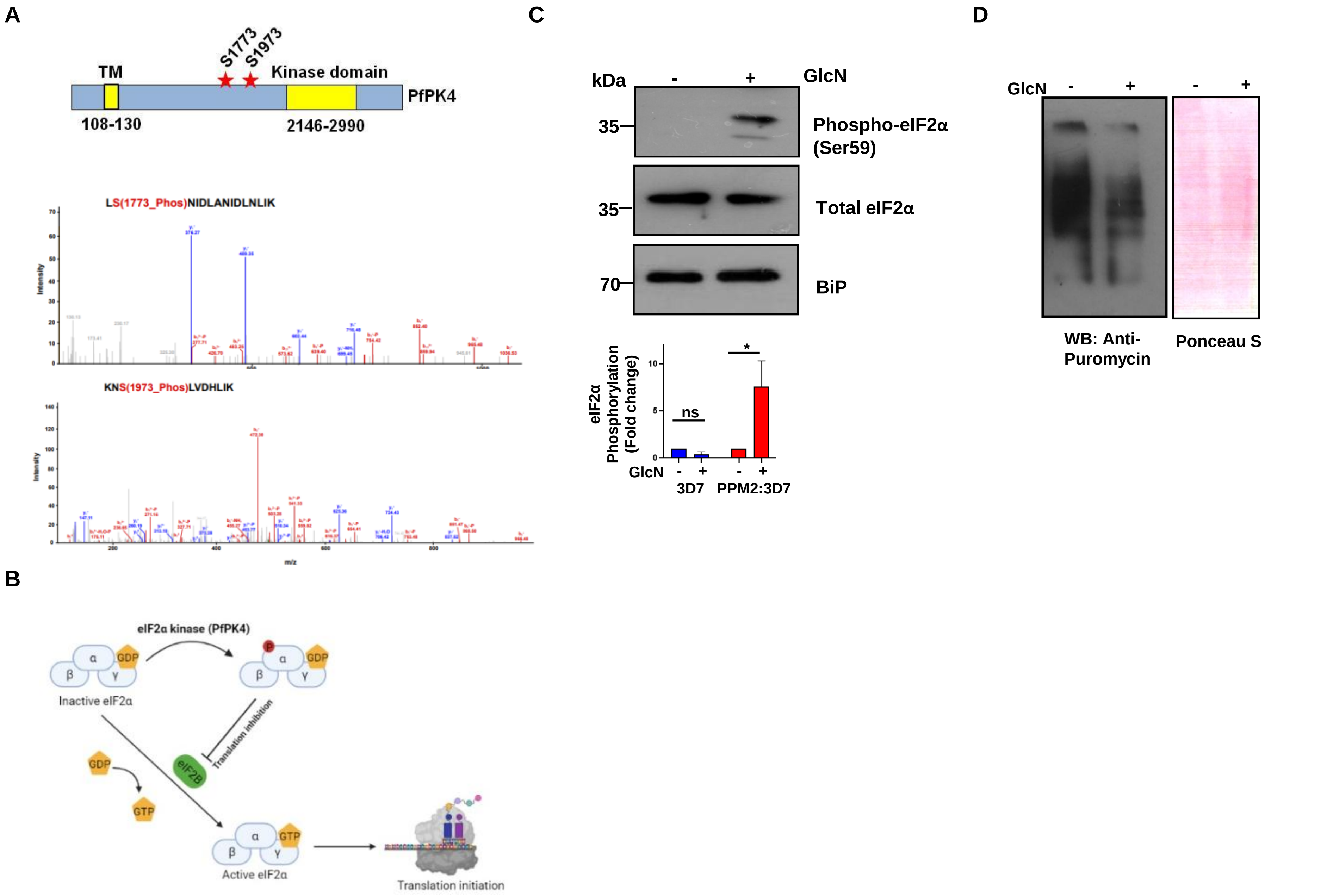
A
TM
S1773
S1973
Kinase domain
PfPK4
108-130
2146-2990
LS(1773_Phos)NIDLANIDLNLIK
Intensity
KNS(1973_Phos)LVDHLIK
m/z
B
eIF2α kinase (PfPK4)
GDP
Inactive eIF2α
Translation inhibition
eIF2B
GTP
Active eIF2α
Translation initiation
C
kDa
GlcN
35
Phospho-eIF2α (Ser59)
Total eIF2α
70
BiP
eIF2α Phosphorylation (Fold change)
ns
*
3D7
PPM2:3D7
D
WB: Anti-Puromycin
Ponceau S

Fig. S5

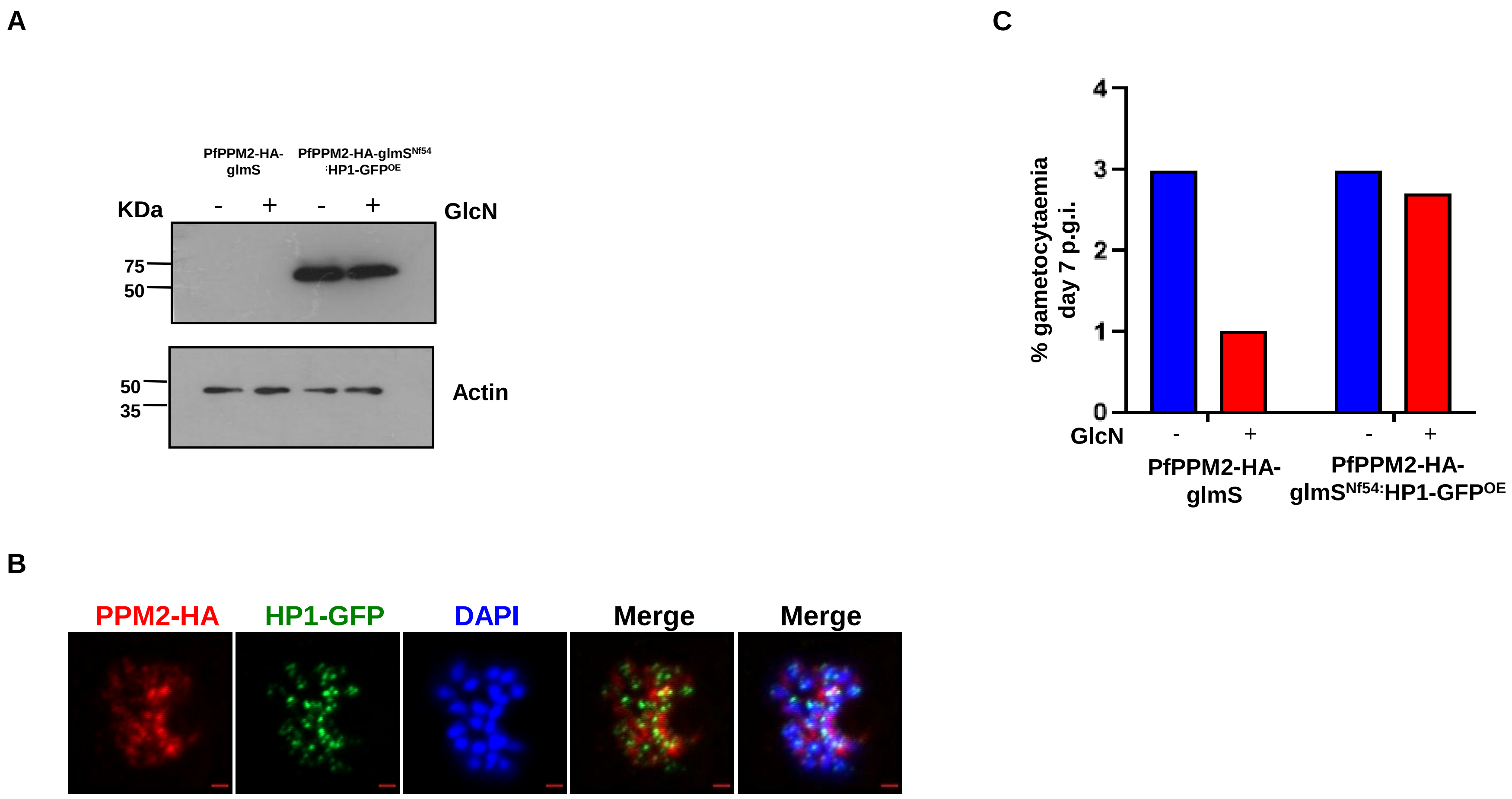
A
PfPPM2-HA-glmS
PfPPM2-HA-glmSNf54:HP1-GFPOE
KDa
- + - +
GlcN
75
50
50
35
Actin
C
% gametocytaemia day 7 p.g.i.
4
3
2
1
0
GlcN
- + - +
PfPPM2-HA-glmS
PfPPM2-HA-glmSNf54:HP1-GFPOE
B
PPM2-HA
HP1-GFP
DAPI
Merge
Merge

Fig. S6

A

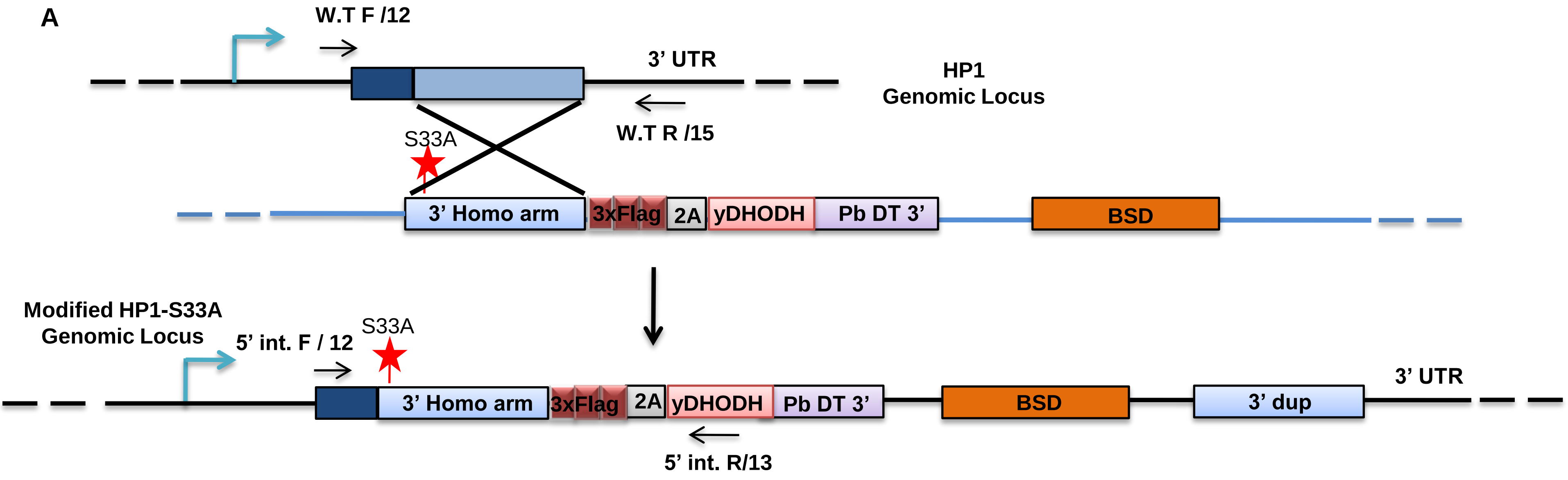


B

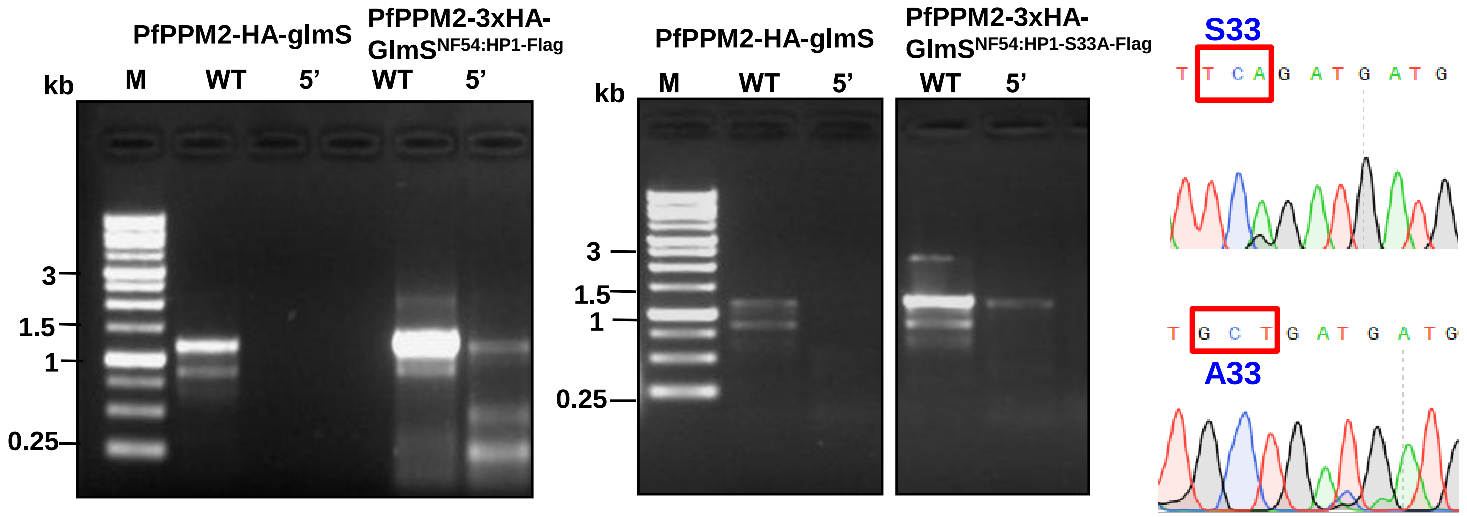


| Locus | Primer set | Expected size |
|---|---|---|
| WT | 12/15 | 1.2 kb |
| 5' integration | 12/13 | 1.1 kb |

Fig. S7

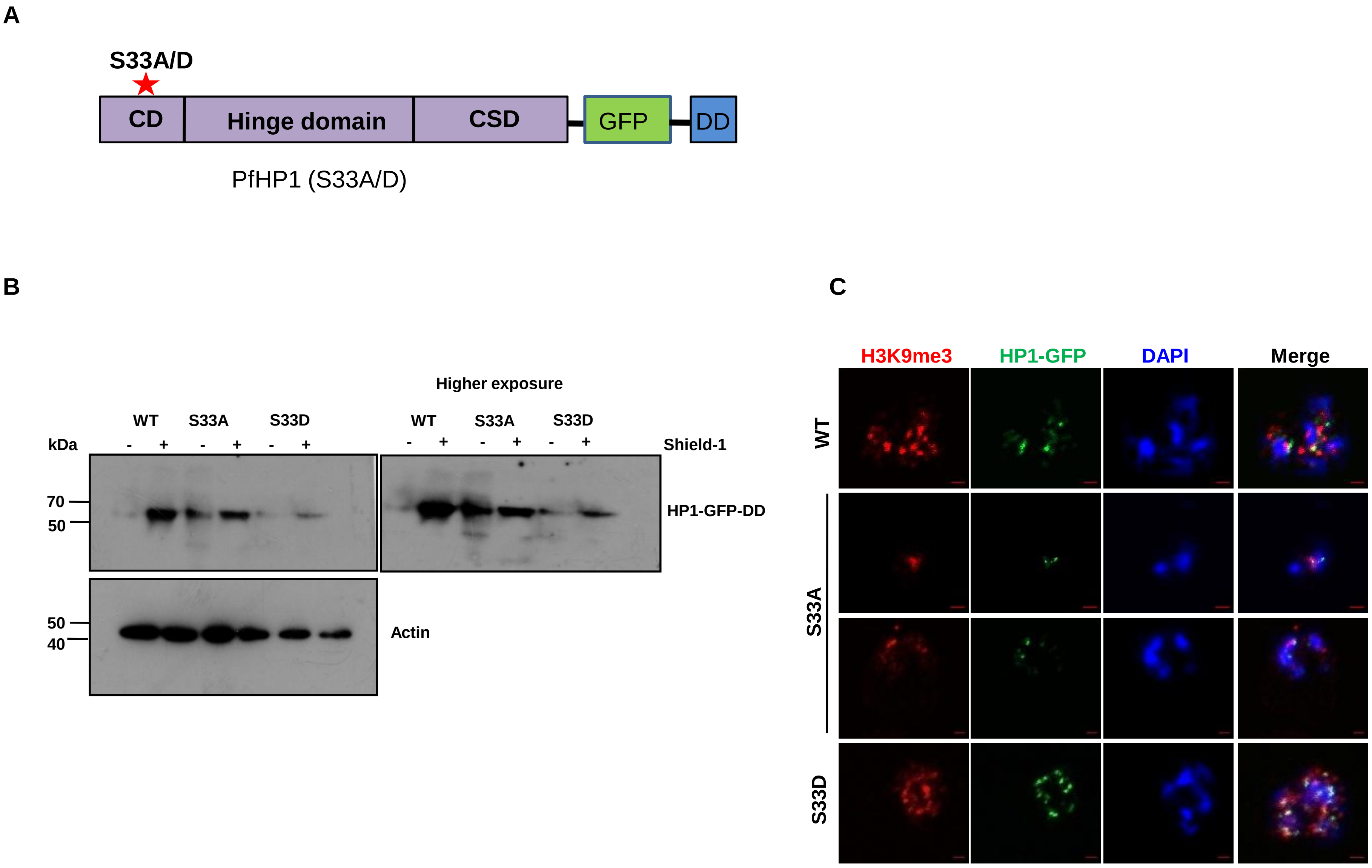
A
S33A/D
CD
Hinge domain
CSD
GFP
DD
PfHP1 (S33A/D)
B
Higher exposure
WT
S33A
S33D
kDa
- + - + - +
Shield-1
70
50
HP1-GFP-DD
50
40
Actin
C
H3K9me3
HP1-GFP
DAPI
Merge
WT
S33A
S33D

Fig. S8

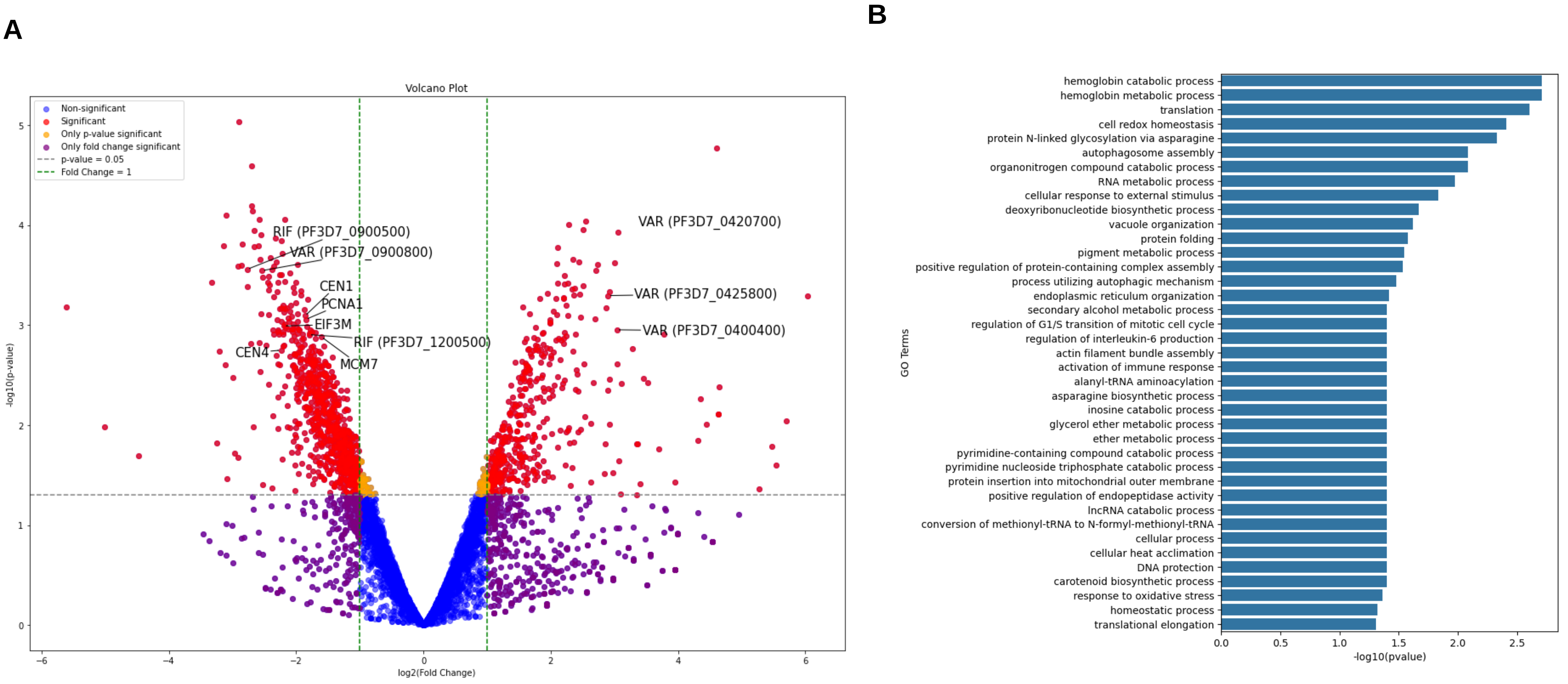
A
Volcano Plot
Non-significant
Significant
Only p-value significant
Only fold change significant
p-value = 0.05
Fold Change = 1
RIF (PF3D7_0900500)
VAR (PF3D7_0900800)
CEN1
PCNA1
EIF3M
RIF (PF3D7_1200500)
CEN4
MCM7
VAR (PF3D7_0420700)
VAR (PF3D7_0425800)
VAR (PF3D7_0400400)
-log10(p-value)
log2(Fold Change)
B
GO Terms
hemoglobin catabolic process
hemoglobin metabolic process
translation
cell redox homeostasis
protein N-linked glycosylation via asparagine
autophagosome assembly
organonitrogen compound catabolic process
RNA metabolic process
cellular response to external stimulus
deoxyribonucleotide biosynthetic process
vacuole organization
protein folding
pigment metabolic process
positive regulation of protein-containing complex assembly
process utilizing autophagic mechanism
endoplasmic reticulum organization
secondary alcohol metabolic process
regulation of G1/S transition of mitotic cell cycle
regulation of interleukin-6 production
actin filament bundle assembly
activation of immune response
alanyl-tRNA aminoacylation
asparagine biosynthetic process
inosine catabolic process
glycerol ether metabolic process
ether metabolic process
pyrimidine-containing compound catabolic process
pyrimidine nucleoside triphosphate catabolic process
protein insertion into mitochondrial outer membrane
positive regulation of endopeptidase activity
lncRNA catabolic process
conversion of methionyl-tRNA to N-formyl-methionyl-tRNA
cellular process
cellular heat acclimation
DNA protection
carotenoid biosynthetic process
response to oxidative stress
homeostatic process
translational elongation
-log10(pvalue)

**Fig. S9**

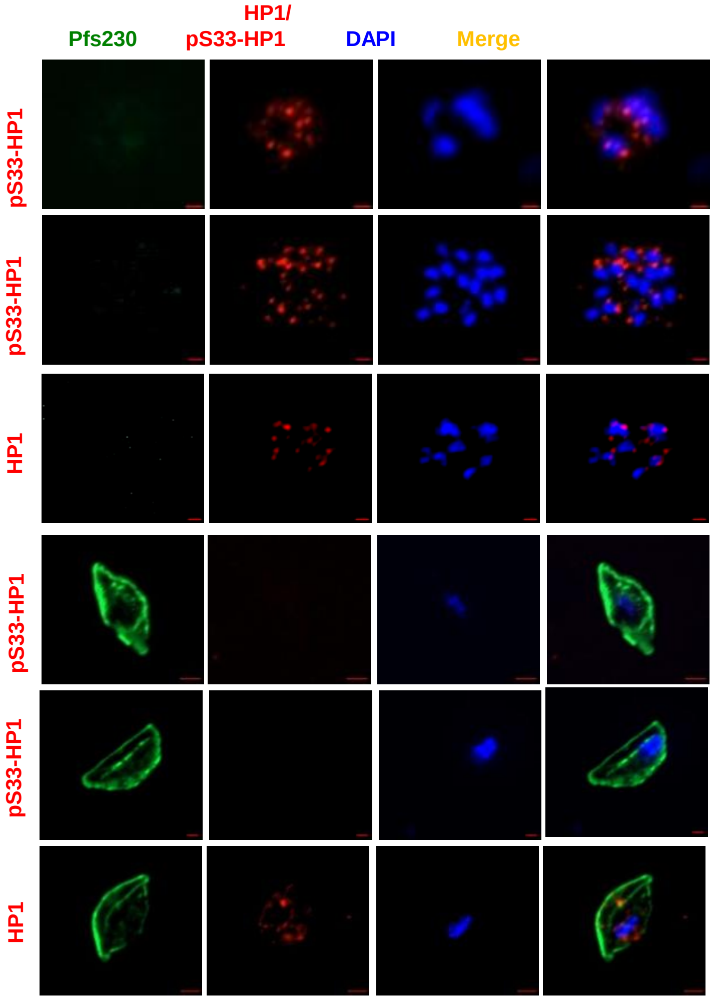

HP1/
Pfs230
pS33-HP1
DAPI
Merge
pS33-HP1
pS33-HP1
HP1
pS33-HP1
pS33-HP1
HP1


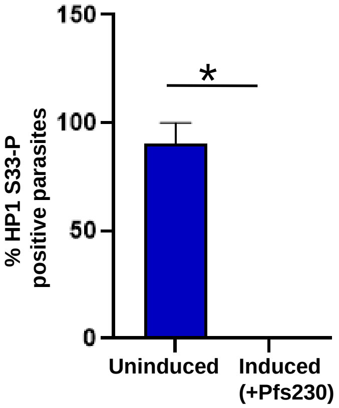

150
100
50
0
*
% HP1 S33-P
positive parasites
Uninduced
Induced
(+Pfs230)


Fig. S10
